# SELENOT regulates endoplasmic reticulum calcium flux via SERCA2 and maintains dopaminergic DAT to protect against attention deficit hyperactivity disorder in mice

**DOI:** 10.1101/2023.11.07.565948

**Authors:** Qing Guo, Zhao-Feng Li, Dong-Yan Hu, Pei-Jun Li, Kai-Nian Wu, Hui-Hui Fan, Jie Deng, Hong-Mei Wu, Xiong Zhang, Jian-Hong Zhu

## Abstract

Attention deficit and hyperactivity disorder (ADHD) is a prevalent developmental disorder. SELENOT is an endoplasmic reticulum-resident selenocysteine-containing protein. We aimed to investigate the role of SELENOT in dopaminergic neurons. Results from *Selenot^fl/fl^;Dat-cre* mice showed that SELENOT deficiency in dopaminergic neurons resulted in ADHD-like behaviors including hyperlocomotion, recognition memory deficit, repetitive movement, and impulsivity. Dopamine metabolism, extrasynaptic dopamine, spontaneous excitatory postsynaptic currents in the striatum and electroencephalogram theta power were enhanced in *Selenot^fl/fl^;Dat-cre* mice, whereas dopaminergic neurons in the substantia nigra were slightly reduced but exhibited normal neuronal firing and little cellular stress. Among dopamine-associated proteins, dopamine transporter (DAT) level was remarkably reduced and monoamine oxidase A increased mildly in the striatum and/or midbrain of *Selenot^fl/fl^;Dat-cre* mice. The ADHD-like phenotype and DAT ablation were corroborated in *Selenot^fl/fl^;Nestin- cre* mice, but not in *Selenot^fl/fl^;Gfap-cre* mice. In vitro overexpression and knockdown analyses and RNA-sequencing data revealed that SELENOT causatively regulated DAT mRNA and protein expression through Ca^2+^ signaling and NURR1. SELENOT maintained cellular Ca^2+^ levels via interaction with endoplasmic reticulum SERCA2, but not IP3Rs and RYRs, as demonstrated by Ca^2+^ imaging, co-immunoprecipitation coupled with mass spectrometry, and colocalization analyses. Treatment with psychostimulants, amphetamine or methylphenidate, rescued the hyperactivity in *Selenot^fl/fl^;Dat-cre* mice. In conclusion, SELENOT in dopaminergic neurons is indispensable to maintain proper dopamine signaling in the midbrain against ADHD.

## Introduction

Midbrain dopaminergic neurons abundantly reside in the substantia nigra pars compacta (SNpc) and ventral tegmental area (VTA). The nigrostriatal projection of dopaminergic neurons from the SNpc to dorsal striatum mainly controls voluntary movement as part of the basal ganglia circuit. The mesolimbic projection of dopaminergic neurons from the VTA to ventral striatum and the mesocortical projection from the VTA to prefrontal cortex regulate multiple functions, including reward, addiction, emotion, and motivation. Striatum-derived postsynaptic dopamine mediates nigrostriatal and corticostriatal transmission in medium spiny neurons bearing dopamine D1 receptor (D1R) and/or dopamine D2 receptor (D2R), thereby controlling movement and some cognitive functions (1, 2). Defects in midbrain dopaminergic neurons have been implicated in multiple neurological and psychiatric disorders such as Parkinson’s disease, Schizophrenia, attention deficit and hyperactivity disorder (ADHD), and drug addiction (3, 4).

Selenoprotein T (SELENOT), a selenocysteine (Sec)-containing and an evolutionary conserved protein, is anchored at the endoplasmic reticulum (ER) membrane through a hydrophobic domain (5, 6). SELENOT possesses a thioredoxin reductase-like activity centered on the Sec residue (7), and is implicated in cytosolic Ca^2+^ maintenance in PC12 cells and SK-N-SH cells, potentially via the Sec-mediated redox regulation (5, 8). The mouse Encyclopedia of DNA Elements Project predicts that SELENOT is preferentially expressed in central nervous system, cerebellum, and kidney (9). SELENOT is known to regulate hormone maturation and secretion in endocrine cells (10), and safeguard dopaminergic neurons in cellular and mouse models of Parkinson’s disease (7, 8). Whole-body *Selenot* deficiency leads to early embryonic lethality in mice (7).

As a prevalent and heterogeneous disorder, ADHD is manifested with predominant symptoms in the domains of inattention, hyperactivity and impulsivity, or combined that mostly occur in children and often extend into adolescence and adulthood. This developmental disorder is highly heritable, associated with at least 14 candidate genes according to genome-wide association or individual gene studies, and prone to environmental and social exposures (11, 12). While the etiology of ADHD is little understood, its pathophysiology has been linked to frontoparietal, dorsal frontostriatal and mesocorticolimbic circuits, and the default mode network and cognitive control network (12). Herein, we developed three lines of *Selenot* conditional knockout mice and demonstrated essential roles of SELENOT in the dopaminergic neurons to prevent ADHD-like phenotypes. The underlying mechanisms comprise a regulatory cascade involving ER-cytosol Ca^2+^ flux, presynaptic dopamine transporter (DAT), and postsynaptic excitability.

## Results

### *Selenot^fl/fl^;Dat-cre* mice display such ADHD-like behaviors as hyperlocomotion, recognition memory deficit, repetitive movement, and impulsivity

Conditional knockout of *Selenot* in dopaminergic neurons (*Selenot^fl/fl^;Dat-cre*) was generated through *Dat* promoter-mediated CRE excision (Figure 1A) and confirmed by co-immunostaining of SELENOT and tyrosine hydroxylase (TH) in the substantia nigra (Figure 1B). The development of *Selenot^fl/fl^;Dat-cre* mice and *Selenot^fl/fl^* control mice was comparable as indicated by 1) the cortex and hippocampus volumes at postnatal day 3 and week 6 (Supplementary Figure 1), and 2) body weight at week 8 (Figure 1C). Results of open field test (Figure 1D) showed that spontaneous locomotion (distanced traveled every 10 min and total distance; Figure 1, E and F), active time, and mean moving speed (Figure 1, G and H) were significantly elevated in *Selenot^fl/fl^;Dat-cre* mice compared to *Selenot^fl/fl^* mice. These two lines of mice exhibited comparable amount of center distance and center time in the open field (Supplementary Figure 2, A and B), suggesting no anxiety in *Selenot^fl/fl^;Dat-cre* mice.

**Figure 1.**
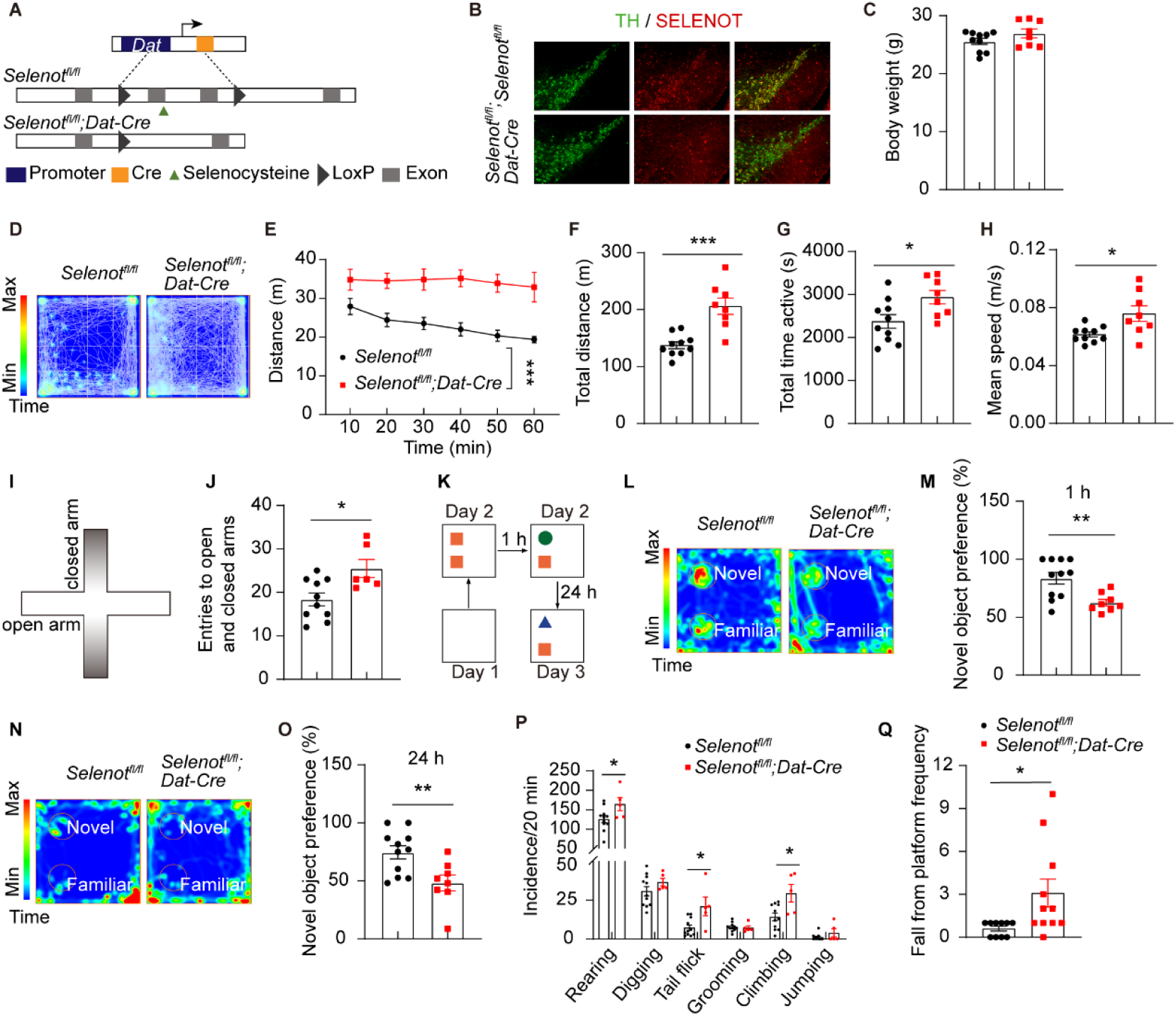
*Selenot^fl/fl^;Dat-cre* mice exhibit hyperlocomotion and impaired memory. (**A**) Schematic diagram of *Dat* promoter-driven excision of *Selenot* exons 2 and 3. (**B**) Immunofluorescence analyses of SELENOT expression in dopaminergic neurons of the substantia nigra in *Selenot^fl/fl^* and *Selenot^fl/fl^;Dat-cre* mice. Green, TH; Red, SELENOT. (**C**) Body weight of *Selenot^fl/fl^*(n = 10) and *Selenot^fl/fl^;Dat-cre* (n = 8) mice aged 8 weeks. (**D**-**H**) Open field test of *Selenot^fl/fl^* (n = 10) and *Selenot^fl/fl^;Dat-cre* (n = 8) mice. Presented are representative activity heatmap (**D**), distance traveled every 10 min (**E**), total distance traveled (**F**), total active time (**G**), and mean speed (**H**). (**I** and **J**) Elevated plus maze test of *Selenot^fl/fl^* (n = 10) and *Selenot^fl/fl^;Dat-cre* (n = 6) mice. Presented are schematic diagram (**I**) and total entries to open and closed arms (**J**). (**K**-**O**) Novel object recognition test of *Selenot^fl/fl^*(n = 11) and *Selenot^fl/fl^;Dat-cre* (n = 8) mice. Presented are schematic diagram (**K**), representative activity heatmap and novel object preference in short-term memory (1 h; **L** and **M**) and long-term memory (24 h; **N** and **O**). (**P**) Stereotyped behaviors of *Selenot^fl/fl^*(n = 11) and *Selenot^fl/fl^;Dat-cre* (n = 5) mice. (**Q**) Fall from platform frequencies of *Selenot^fl/fl^* (n = 10) and *Selenot^fl/fl^;Dat-cre* (n = 11) mice in cliff avoidance reaction test. Data are presented as means ± SEM and analyzed by two-way repeated measures ANOVA for (**E**) and two-tailed unpaired *t*-test for (**C**, and **F**-**Q**). \**P* < 0.05; \*\**P* < 0.01; \*\*\**P* < 0.001. TH, tyrosine hydroxylase.

Results of elevated plus maze test (Figure 1I) confirmed hyperlocomotion (increased entries to arms; Figure 1J) and no anxiety (increased entries to open arms but comparable amount of time in open arms; Supplementary Figure 2, C and D) in *Selenot^fl/fl^;Dat-cre* mice.

Compared to *Selenot^fl/fl^* mice, *Selenot^fl/fl^;Dat-cre* mice showed 1) impairment in short-term (1 hour; Figure 1, L and M) and long-term memory (24 hours; Figure 1, N and O) according to novel object recognition test (Figure 1K), 2) repetitive behaviors in rearing, tail flick, and climbing (Figure 1P), and 3) enhanced impulsive behavior based on cliff avoidance reaction test (Figure 1Q). In contrast, *Selenot^fl/fl^;Dat-cre* mice appeared normal in motor coordination according to rotarod test (Supplementary Figure 2, E and F), spatial memory according to Y-maze test (Supplementary Figure 2G), frequencies of self-grooming, jumping, and digging (Figure 1P), and social interaction according to three chamber sociability test including social preference (Supplementary Figure 2, H-K) and social recognition (Supplementary Figure 2, L-O).

### Impaired dopamine metabolism and reuptake and enhanced postsynaptic excitability in the striatum of *Selenot^fl/fl^;Dat-cre* mice

To gain insight into the behavioral changes, key striatal neurotransmitters [norepinephrine, γ- aminobutyric acid (GABA), acetylcholine, 5-hydroxytryptamine, and dopamine (DA)] and DA metabolites [3,4-dihydroxyphenylacetic acid (DOPAC), homovanillic acid (HVA), and 3- methoxytyramine (3-MT)] were assessed in the whole striatum. The results showed that levels of these chemicals were not significantly different between *Selenot^fl/fl^;Dat-cre* and *Selenot^fl/fl^* mice (Figure 2, A-E). However, ratios of 3-MT/DA, HVA/DA and DOPAC/DA were increased in *Selenot^fl/fl^;Dat-cre* mice (Figure 2F), suggesting an accelerated dopamine metabolism. Next, results using striatal microdialyzed fluids from live mice (Figure 2G) showed that the extrasynaptic dopamine level was significantly higher in *Selenot^fl/fl^;Dat-cre* mice than in *Selenot^fl/fl^* mice (Figure 2H). These indicate that dopamine metabolism was impaired and a high level of dopamine was retained in the synaptic gap in the striatum of *Selenot^fl/fl^;Dat-cre* mice. In addition, levels of glutamic acid (Figure 2I), but not 5- hydroxyindoleacetic acid (Figure 2J; a 5-hydroxytryptamine metabolite), were increased in the striatum of *Selenot^fl/fl^;Dat-cre* mice.

**Figure 2.**
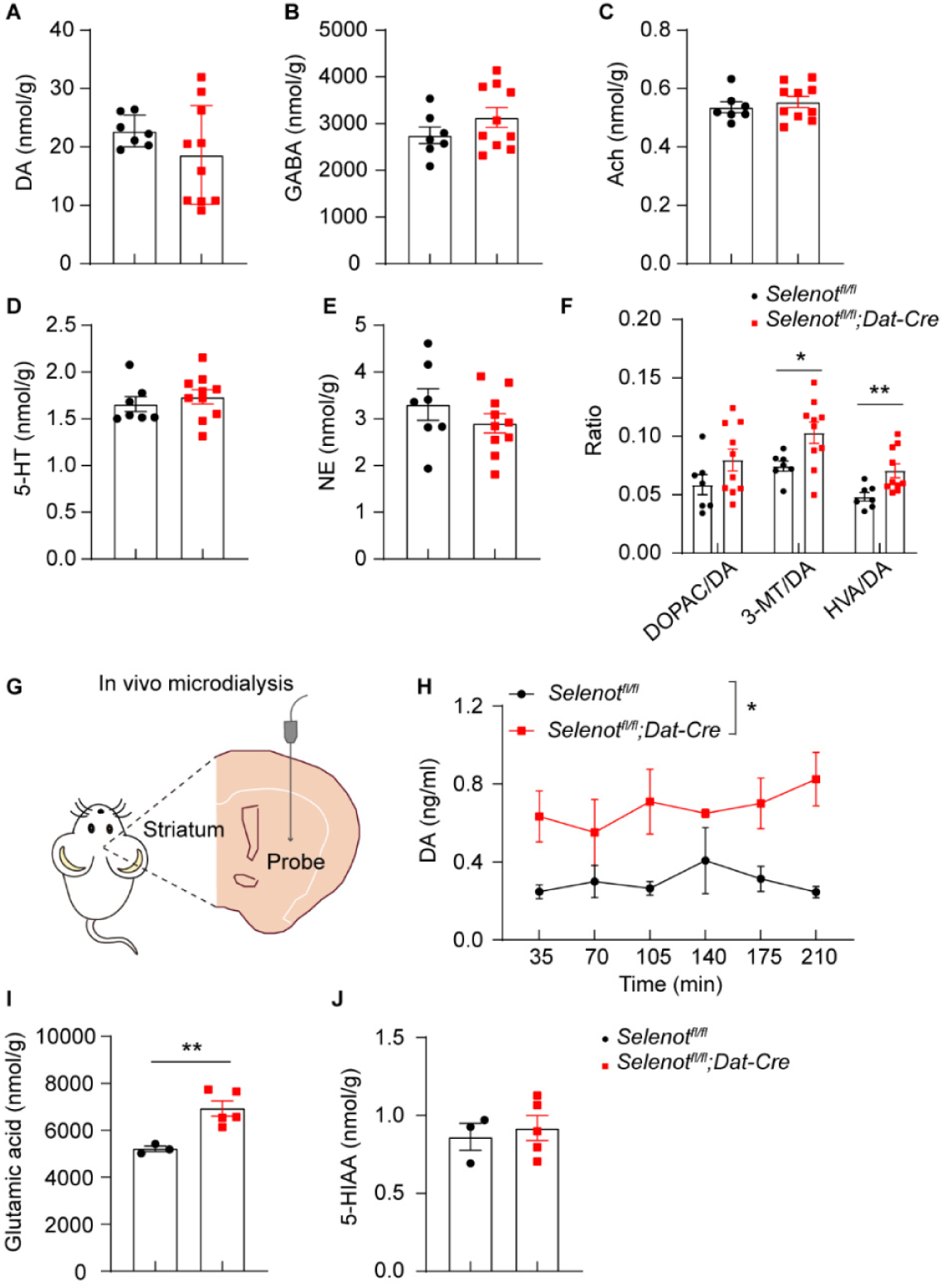
Impaired DA metabolism and reuptake in *Selenot^fl/fl^;Dat-cre* mice. (**A**-**E**) HPLC analyses of striatal neurotransmitters, including DA (**A**), GABA (**B**), Ach (**C**), 5-HT (**D**) and NE (**E**). (**F**) Ratio of DA metabolites to DA. n = 7 *Selenot^fl/fl^* mice and n = 10 *Selenot^fl/fl^;Dat- cre* mice. (**G** and **H**) DA levels measured in the striatum by microdialysis in freely moving *Selenot^fl/fl^*(n = 3) and *Selenot^fl/fl^;Dat-cre* (n = 4) mice. Presented are schematic diagram (**G**) and HPLC analyzed DA levels (**H**). (**I** and **J**) HPLC analyses of striatal glutamate (**I**) and 5- HIAA (**J**) levels. Data are presented as means ± SEM and analyzed by two-way repeated measures ANOVA for (**H**) and two-tailed unpaired *t*-test for (**A**-**F**, **I**, and **J**). \**P* < 0.05; \*\**P* < 0.01. 3-MT, 3-methoxytyramine; 5-HT, 5-hydroxytryptamine; 5-HIAA, 5- hydroxyindoleacetic acid; Ach, acetylcholine; DA, dopamine; DOPAC, 3,4- dihydroxyphenylacetic acid; GABA, γ-aminobutyric acid; HVA, homovanillic acid; NE, norepinephrine.

We next determined whether *Selenot* deficiency affects neural activity by using cell patch clamp recording. Dopaminergic neurons were evidenced by the co-immunostaining of biocytin and TH (Figure 3A). The threshold and amplitude of action potential remained unchanged in the substantia nigra of *Selenot^fl/fl^;Dat-cre* mice (Figure 3, B-D). The pacemaker frequency was also not changed in the mice (Supplementary Figure 3, A and B). Analyses of spontaneous excitatory postsynaptic currents (sEPSC) showed that the frequency, but not the amplitude, in medium spiny neurons was significantly enhanced in the striatum of *Selenot^fl/fl^;Dat-cre* mice (Figure 3, E-G). Altogether, these results implicate extrasynaptic dopamine upregulation in the striatum of *Selenot^fl/fl^;Dat-cre* mice. Further electroencephalogram (EEG) analyses (Figure 3, H-J) showed that theta (3-8 Hz) power elevated significantly (Figure 3K), while EEG power of higher frequency (> 8 Hz) appeared normal (Supplementary Figure 3, C and D) in occipital regions of *Selenot^fl/fl^;Dat-cre* mice.

**Figure 3.**
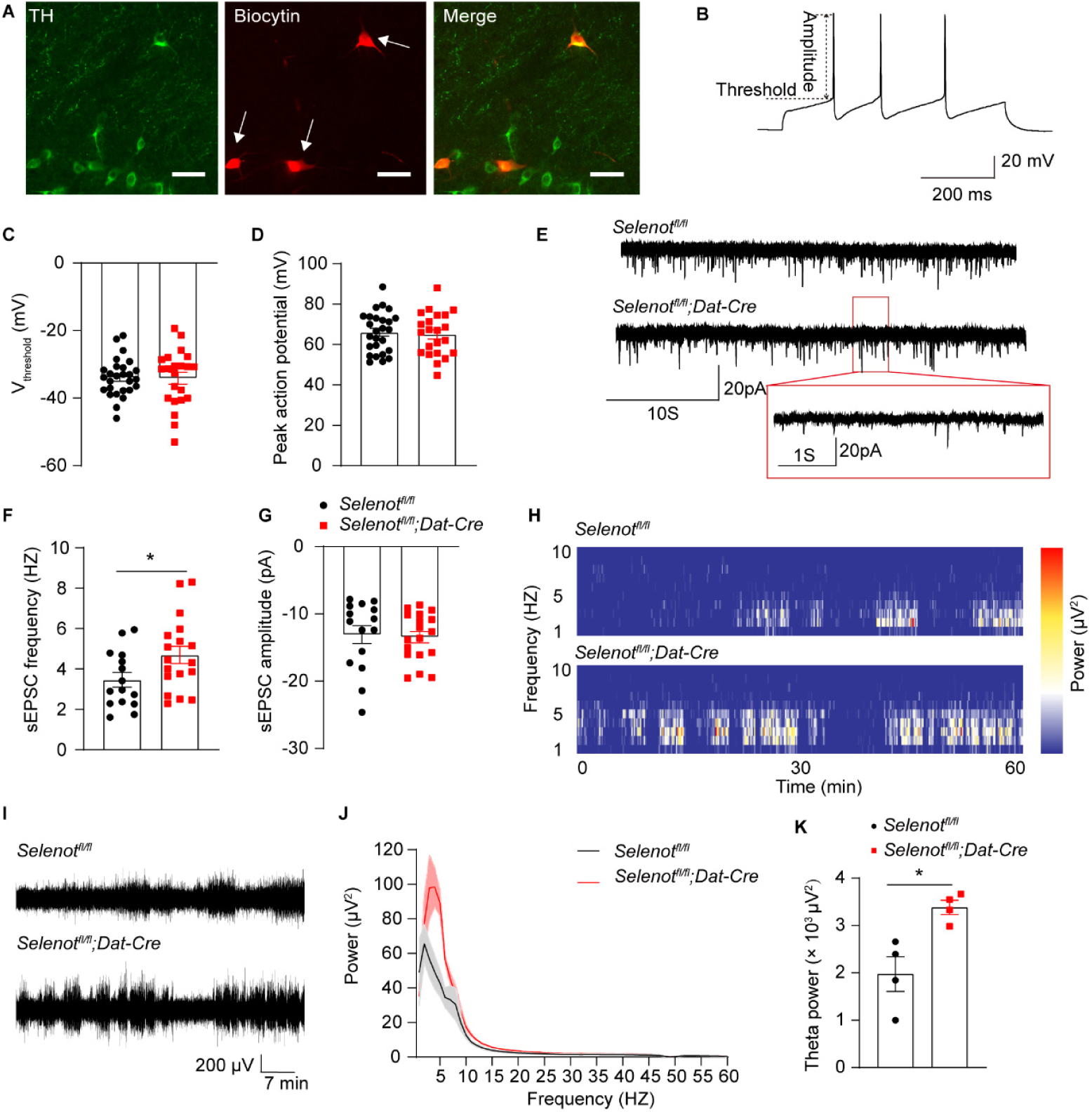
*Selenot^fl/fl^;Dat-cre* mice display normal dopaminergic neuron firing but elevated postsynaptic excitability and EEG theta power. (**A**) Representative images showing that the recorded neurons (white arrows) were positive for TH (green) on a polyoxymethylene-fixed brain slice after patching. Biocytin (red) was injected into cells by whole cell patch clamp. Scale bar, 50 μm. (**B**-**D**) Action potential of dopaminergic neurons. Presented are representative action potential recorded by current clamp (**B**), and quantified thresholds (**C**) and amplitudes (**D**) of action potentials. n = 26-27 neurons recorded from 8 *Selenot^fl/fl^* mice and n = 22 neurons recorded from 8 *Selenot^fl/fl^;Dat-cre* mice. (**E**-**G**) sEPSC in the striatum. Presented are representative recordings of sEPSC (**E**), and quantified frequencies (**F**) and amplitudes (**G**) of striatal medium spiny neurons. n = 15 neurons recorded from 4 *Selenot^fl/fl^*mice and n = 18 neurons recorded from 3 *Selenot^fl/fl^;Dat-cre* mice. (**H**-**K**) EEG analysis. Presented are representative spectrogram (**H**) and traces (**I**), and mean EEG power spectral density (**J**) and theta power (**K**) in 60 min. n = 4 *Selenot^fl/fl^* mice and n = 4 *Selenot^fl/fl^;Dat-cre* mice. Data are presented as means ± SEM and analyzed by two-tailed unpaired *t*-test. \**P* < 0.05. EEG, electroencephalogram; sEPSC, spontaneous excitatory postsynaptic currents; TH, tyrosine hydroxylase; V_threshold_, threshold of action potential.

### *Selenot^fl/fl^;Dat-cre* mice exhibit remarkably reduced DAT expression and mildly elevated MAO-A expression, but no induction in midbrain cellular stress

To better understand why the striatal extrasynaptic dopamine level was elevated, we examined levels of proteins in association with dopamine synthesis (i.e., TH), signaling [i.e., vesicular monoamine transporter 2 (VMAT2), D1R, and D2R], reuptake (i.e., DAT), and metabolism [i.e., catechol-O-methyltransferase (COMT), monoamine oxidase A (MAO-A), and MAO-B] by Western blot. Results showed that DAT level was remarkably reduced (by 57% and 79%, respectively) but MAO-A increased in both dorsal and ventral striatum of *Selenot^fl/fl^;Dat-cre* mice. Levels of other proteins including TH were comparable between *Selenot^fl/fl^;Dat-cre* and *Selenot^fl/fl^* mice (Figure 4, A and B). Such findings of DAT and TH were corroborated with striatal immunohistochemical analyses (Figure 4, C-F). In line with that in the striatum, DAT level was drastically reduced in the substantia nigra (by 61%) and in the VTA (by 85%) of *Selenot^fl/fl^;Dat-cre* mice; however, TH level was mildly reduced in these two regions as suggested by the Western blot analyses (Figure 4, G and H). Further immunostaining analyses demonstrated that quantity of dopaminergic neurons was mildly reduced in the midbrain of *Selenot^fl/fl^;Dat-cre* mice (Figure 4, I and J); however, while cellular DAT expression was indeed reduced, cellular TH expression remained unchanged in the dopaminergic neurons of *Selenot^fl/fl^;Dat-cre* mice (Figure 4, K and L). In addition, DAT expression did not differ in the substantia nigra or striatum between *Selenot^fl/fl^* mice and *Dat-cre* mice (Supplementary Figure 4, A-D), precluding potential interference of *Dat-cre* on DAT expression. The substantia nigra of *Selenot^fl/fl^;Dat-cre* mice were not prone to apoptosis as determined by Western blot analyses of caspase-3, cleaved caspase-3, and B-cell lymphoma-2 (BCL-2), and did not display ER stress as indicated by analyses of binding-immunoglobulin protein (BIP) and CCAAT-enhancer-binding protein homologous protein (CHOP) (Supplementary Figure 4, E and F).

**Figure 4.**
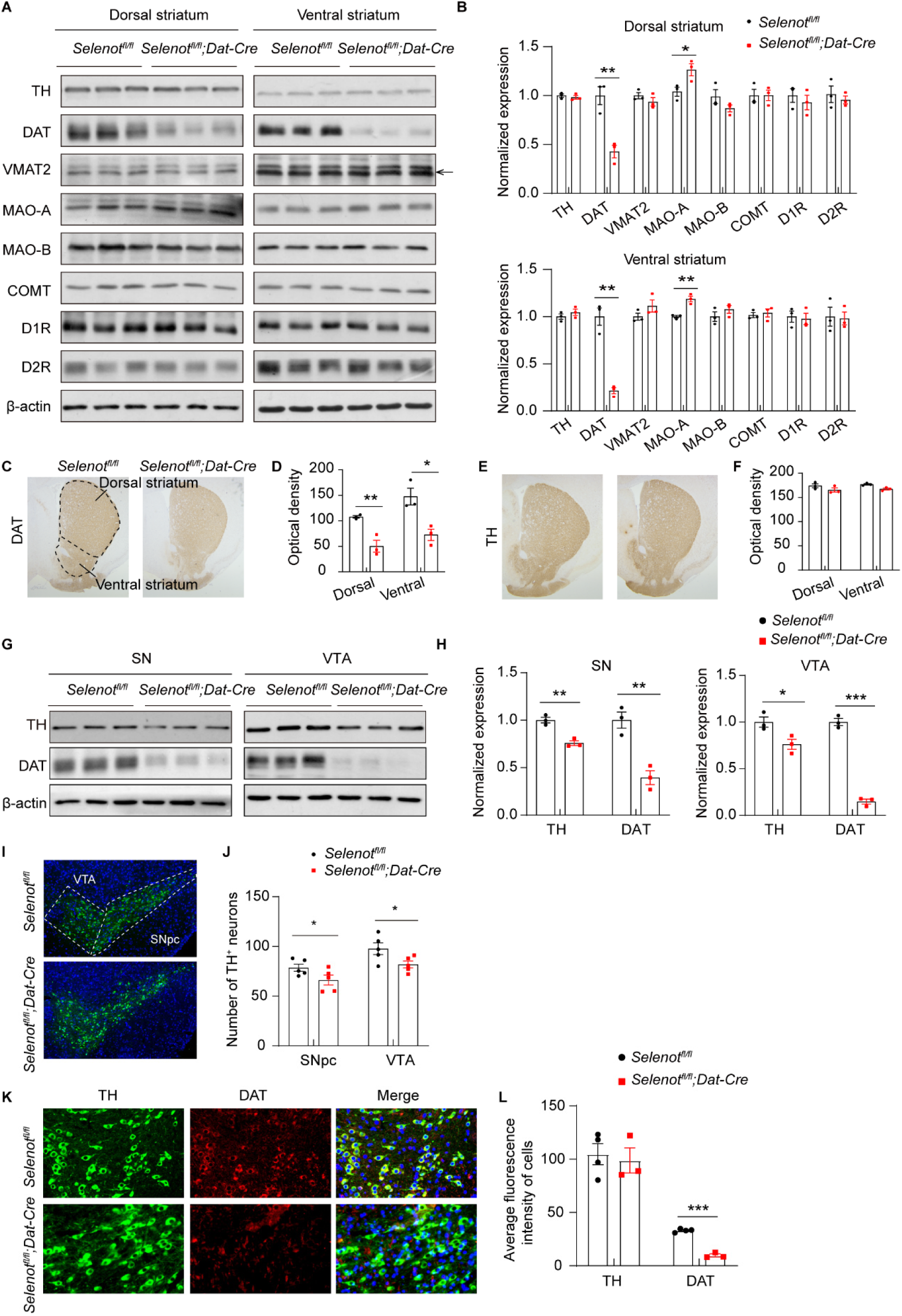
Altered expression of dopamine signaling associated proteins in *Selenot^fl/fl^;Dat- cre* mice. (**A** and **B**) Western blot analyses (**A**) and quantifications (**B**) of the associated proteins in the dorsal and ventral striatum. n = 3 *Selenot^fl/fl^* mice and n = 3 *Selenot^fl/fl^;Dat-cre* mice. (**C**-**F**) Immunohistochemical analyses (**C**, **E**) and optical density quantifications (**D**, **F**) of DAT and TH in the striatum. n = 3 *Selenot^fl/fl^* mice and n = 3 *Selenot^fl/fl^;Dat-cre* mice. (**G** and **H**) Western blot analyses (**G**) and quantifications (**H**) of TH and DAT in the midbrain. n = 3 *Selenot^fl/fl^*mice and n = 3 *Selenot^fl/fl^;Dat-cre* mice. (**I** and **J**) Immunofluorescence (**I**) and counts (**J**) of dopaminergic neurons in the midbrain. n = 5 *Selenot^fl/fl^* mice and n = 5 *Selenot^fl/fl^;Dat-cre* mice. (**K** and **L**) Immunofluorescence (**K**) and cellular quantifications (**L**) of TH and DAT expression in dopaminergic neurons. n = 4 *Selenot^fl/fl^* mice and n = 3 *Selenot^fl/fl^;Dat-cre* mice. Quantifications are normalized to β-actin for Western blot. Each quantification was averaged from 2-3 consecutive slices for immunostaining. Data are presented as means ± SEM and analyzed by two-tailed unpaired *t*-test. \**P* < 0.05; \*\**P* < 0.01; \*\*\**P* < 0.001. COMT, catechol-O-methyltransferase; DAT, dopamine transporter; D1R, dopamine D1 receptor; D2R, dopamine D2 receptor; MAO-A, monoamine oxidase A; MAO- B, monoamine oxidase B; SN, substantia nigra; SNpc, substantia nigra pars compacta; TH, tyrosine hydroxylase; VMAT2, vesicular monoamine transporter 2; VTA, ventral tegmental area.

Together, these results indicate that 1) the accelerated dopamine metabolism is likely associated with the enhanced MAO-A expression; 2) the elevation of extrasynaptic dopamine level corresponds with the DAT reduction and the subsequent impairment in dopamine reuptake; 3) although dopaminergic neurons slightly decreased in the midbrain, the striatal innervations are not affected; 4) the decrease of dopaminergic neurons is not due to cellular stress, but likely resulted from proliferation arrest of progenitors during embryonic development of *Selenot^fl/fl^;Dat-cre* mice given that SELENOT can promote G1-to-S cell cycle transition (8).

### *Selenot* deficiency in whole brain, but not in astrocytes, retains ADHD-like behaviors and reduces DAT expression

To understand whether such a role of SELENOT is specific in dopaminergic neurons, we additionally constructed *Selenot^fl/fl^;Nestin-cre* and *Selenot^fl/fl^;Gfap-cre* mice to respectively knock out *Selenot* expression in the whole brain and astrocytes. The knockouts were confirmed by Western blot and/or immunohistochemical analyses (Supplementary Figure 5, A-C, and Supplementary Figure 6, A and B). Although *Selenot^fl/fl^;Gfap-cre* mice grew normally, body weight of *Selenot^fl/fl^;Nestin-cre* mice was lower than that of *Selenot^fl/fl^* mice at 8 weeks of age (Supplementary Figure 5D) despite no signs of premature cell death. The mice were assessed for the ADHD-like behaviors and DAT expression.

In open field test (Figure 5A), *Selenot^fl/fl^;Nestin-cre* mice exhibited higher spontaneous locomotion (distanced traveled every 10 min and total distance) and active time (Figure 5, B-D), but normal mean speed (Figure 5E), center distance and center time (Supplementary Figure 5, E and F), in comparison to *Selenot^fl/fl^*mice. In elevated plus maze test, *Selenot^fl/fl^;Nestin-cre* mice showed increased entries to arms (Figure 5F), but normal percentages of open time and of open entries (Supplementary Figure 5, G and H). These results suggest hyperlocomotion and no anxiety in *Selenot^fl/fl^;Nestin-cre* mice. By contrast, *Selenot^fl/fl^;Nestin-cre* mice displayed no impairment in 1) motor coordination according to latency to fall rotarod test (Supplementary Figure 5, I and J), 2) in short-term and long-term recognition memory as suggested by novel object recognition tests (Figure 5, G-J), and 3) social interaction, including social preference (Supplementary Figure 5, K-M) and social recognition (Supplementary Figure 5, N-P) based on three-chamber sociability test.

**Figure 5.**
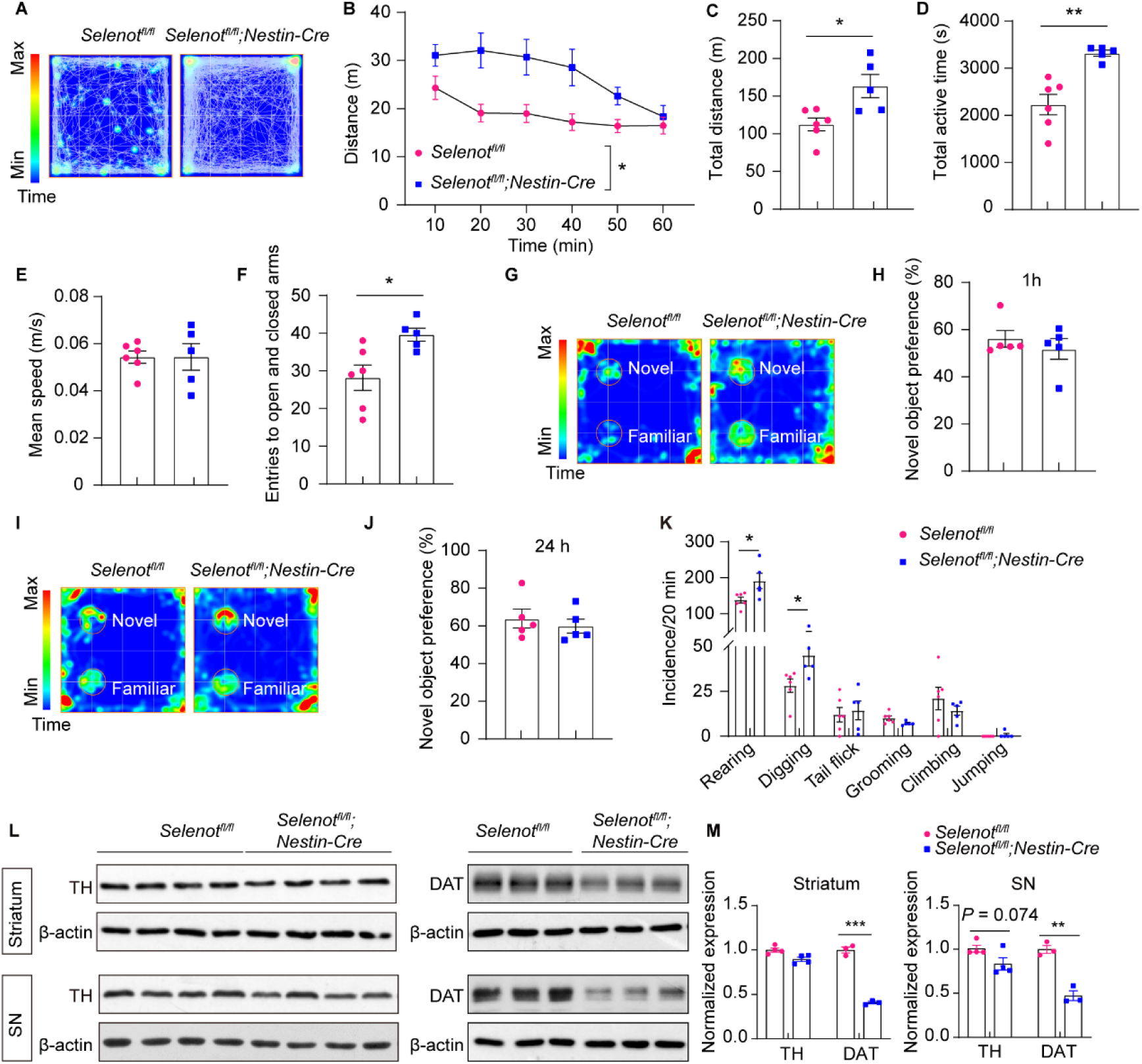
*Selenot^fl/fl^;Nestin-cre* mice exhibit hyperactive behaviors and reduced DAT expression. (**A**-**E**) Open field test of *Selenot^fl/fl^* (n = 6) and *Selenot^fl/fl^;Nestin-cre* (n = 5) mice. Presented are representative activity heatmap (**A**), distance traveled every 10 min (**B**), total distance traveled (**C**), total active time (**D**), and mean speed (**E**). (**F**) Elevated plus maze test of *Selenot^fl/fl^* (n = 6) and *Selenot^fl/fl^;Nestin-cre* (n = 5) mice. Presented are total entries to open and closed arms. (**G**-**J**) Novel object recognition test of *Selenot^fl/fl^* (n = 5) and *Selenot^fl/fl^;Nestin-cre* (n = 5) mice. Presented are representative activity heatmap and novel object preference in short-term memory (1 h; **G** and **H**) and long-term memory (24 h; **I** and **J**). (**K**) Stereotyped behaviors of *Selenot^fl/fl^* (n = 6) and *Selenot^fl/fl^;Nestin-cre* (n = 5) mice. (**L** and **M**) Western blot analysis (**L**) and quantifications (**M**) of TH and DAT in the total striatum and SN. Quantifications are normalized to β-actin. n = 3-4 *Selenot^fl/fl^*mice and n = 3-4 *Selenot^fl/fl^;Nestin-cre* mice. Data are presented as means ± SEM and analyzed by two-way repeated measures ANOVA for (**B**) and two-tailed unpaired *t*-test for (**C**-**M**). \**P* < 0.05; \*\**P* < 0.01; \*\*\**P* < 0.001. DAT, dopamine transporter; SN, substantia nigra; TH, tyrosine hydroxylase.

Furthermore, *Selenot^fl/fl^;Nestin-cre* mice displayed some repetitive behaviors such as rearing and digging, but not others including tail flick, self-grooming, climbing and jumping (Figure 5K). Similar to *Selenot^fl/fl^;Dat-cre* mice, *Selenot^fl/fl^;Nestin-cre* mice exhibited remarkable reduction of DAT levels in the striatum and substantia nigra, and slight reduction of TH levels (*P* = 0.074) in the substantia nigra, but not in the striatum (Figure 5, L and M). Altogether, these results suggest that neurological consequences of *Selenot* deficiency in the whole brain are reminiscent of those in dopaminergic neurons, displaying ADHD-like phenotypes with a marked reduction in DAT expression.

Next, *Selenot^fl/fl^;Gfap-cre* mice were assessed with open field and repetitive behavior tests. Compared to *Selenot^fl/fl^*mice, *Selenot^fl/fl^;Gfap-cre* mice had no signs of 1) hyperactivity as indicated by locomotion, active time, and mean speed in open field test (Supplementary Figure 6, C-G), 2) anxiety as evidenced by normal percentages in center distance and center time in open field test (Supplementary Figure 6, H and I), 3) repetitive behaviors according to rearing, tail flick, self-grooming, climbing and jumping results (Supplementary Figure 6J), and 4) changes in DAT and TH protein expression in the striatum and substantia nigra based on Western blot analyses (Supplementary Figure 6, K and L). Interestingly, the digging frequency was slightly decreased (*P* < 0.05) in the *Selenot^fl/fl^;Gfap-cre* mice (Supplementary Figure 6J). Taken together, these results suggest that *Selenot* deficiency in astrocytes does not lead to ADHD.

### SELENOT directly regulates DAT expression via the transcription factor NURR1

We further explored the link between SELENOT and DAT. In agreement with the in vivo results, knockdown of *SELENOT* led to reduced DAT protein levels in HEK293 cells, while conversely overexpression of *SELENOT* increased DAT protein levels (Figure 6, A and B). Such SELENOT-mediated regulations on *DAT* expression were also observed at transcriptional level (Figure 6C). RNA-seq was then performed using the substantia nigra of *Selenot^fl/fl^*and *Selenot^fl/fl^;Dat-cre* mice to explore the underlying mechanisms. Results of RNA-seq showed a total of 320 differentially expressed genes (DEGs; including 156 upregulated and 164 downregulated; Figure 6D), which however did not contain the 25 selenoprotein genes (Supplementary Figure 7A). Among the dopaminergic signaling-related genes, *Dat*/*Slc6a3* and *Nurr1*/*Nr4a2* were found to be downregulated in the substantia nigra of *Selenot^fl/fl^;Dat-cre* mice (Figure 6E). The *Nurr1*/*Nr4a2* was among the 12 ADHD-associated DEGs revealed in the mice (Figure 6F). The downregulations of *Dat* and *Nurr1* were validated by qPCR analysis using the substantia nigra of *Selenot^fl/fl^* and *Selenot^fl/fl^;Dat- cre* mice (Figure 6G). While such transcriptional change of *Dat* was matched with the earlier Western blot results, we examined *Nurrl* expression by Western blot, which is a known transcription factor for DAT (13). Indeed, NURR1 protein level was significantly reduced in the substantia nigra of *Selenot^fl/fl^;Dat-cre* mice (Figure 6, H and I). Results of siRNA knockdown validated that NURR1 could regulate the transcriptional level of *DAT* (Figure 6J). In accordance with the *DAT* results, knockdown and overexpression of *SELENOT* respectively upregulated and downregulated the protein and mRNA expression of *NURR1* (Figure 6, K-M). Next, we further demonstrated that siRNA knockdown of *NURR1* expression blunted the SELENOT overexpression-induced upregulation of *DAT* level (Figure 6N).

**Figure 6.**
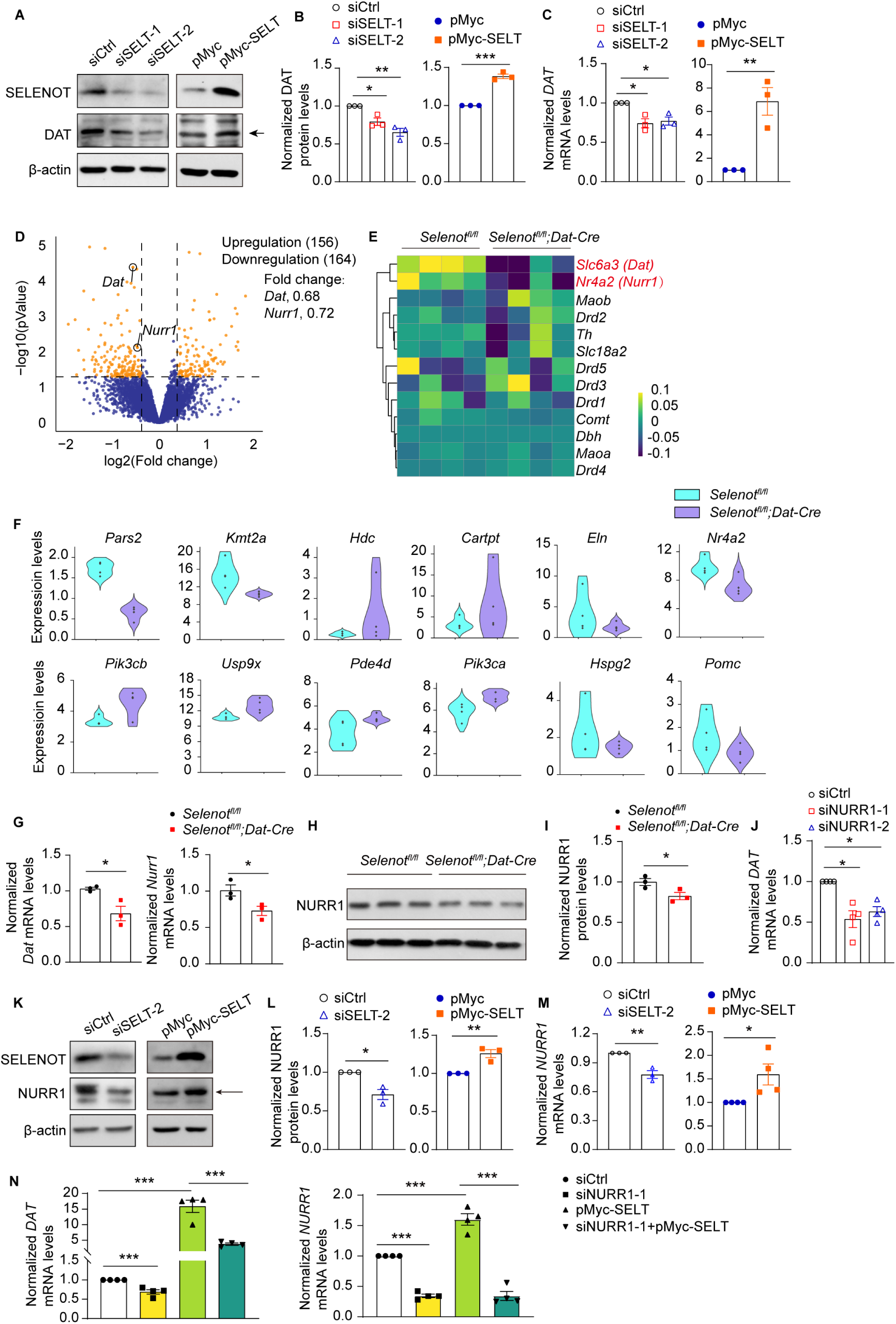
RNA sequencing analysis and regulation of SELENOT on DAT and NURR1 expression. (**A**-**C**) Western blot analyses (**A**) and quantifications (**B**), and qPCR analyses (**C**) for DAT protein and mRNA levels in HEK293 cells transfected with siSELT-1, siSELT-2 or siCtrl, or pMyc-SELT or pMyc (48 h for qPCR and 72 h for Western blot). n = 3 independent experiments. (**D**) Volcano plot of DEGs in the substantia nigra of *Selenot^fl/fl^;Dat-cre* mice comparing to *Selenot^fl/fl^* mice. Fold change > 1.2, *P* < 0.05. (**E**) Heatmap of row-normalized RNA-seq results of dopaminergic signaling related genes in the substantia nigra of *Selenot^fl/fl^*and *Selenot^fl/fl^;Dat-cre mice.* n = 4. (**F**) Violin plots of RNA-seq results of ADHD candidate genes by DisGenet*. P* < 0.05. (**G**) qPCR validation of *Dat* and *Nurr1* mRNA expression in the substantia nigra of *Selenot^fl/fl^* (n = 3) and *Selenot^fl/fl^;Dat-cre* (n = 3) mice. (**H** and **I**) Western blot analysis (**H**) and quantifications (**I**) of NURR1 protein levels in the substantia nigra of *Selenot^fl/fl^*(n = 3) and *Selenot^fl/fl^;Dat-cre* (n = 3) mice. (**J**) Validation of the transcriptional regulation of DAT expression by NURR1. HEK293 cells were transfected with siNURR1-1, siNURR1-2 or siCtrl for 48 h. n = 4 independent experiments. (**K**-**M**) Western blot analyses (**K**) and quantifications (**L**), and qPCR analyses (**M**) for NURR1 protein and mRNA levels in HEK293 cells transfected with siSELT-2 or siCtrl, or pMyc-SELT or pMyc (48 h for qPCR and 72 h for Western blot). n = 3-4 independent experiments. (**N**) qPCR analyses for the effect of NURR1 on the SELENOT-induced *DAT* expression. Cells were pre-transfected with siNURR1 or siCtrl for 24 h, followed by transfection with pMyc-SELT or pMyc for 48 h. n = 4 independent experiments. Quantifications are normalized to β-actin for qPCR and Western blot. Data are presented as means ± SEM and analyzed by one-way ANOVA followed by Dunnett’s post-hoc test for (left panels of **B** and **C**, and **J**), factorial ANOVA for (**N**), and two-tailed unpaired *t*-test for other comparisons. \**P* < 0.05; \*\**P* < 0.01; \*\*\**P* < 0.001. DAT, dopamine transporter; DEG, differentially expressed gene; NURR1, nuclear receptor-related 1; pMyc, pCMV-Myc empty vector; pMyc-SELT, Myc-tagged SELENOT vector; siCtrl, scramble siRNA; siNURR1, *NURR1* siRNA; siSELT, *SELENOT* siRNA.

### SELENOT modulates ER-cytosol Ca^2+^ flux via SERCA channel, thereby regulating NURR1 and DAT expression

Gene Ontology analysis of the DEGs disclosed biological processes of dopamine biosynthetic process, regulation of dopamine metabolic process, and neurotransmitter secretion.

Meanwhile, significant enrichments were exhibited in Ca^2+^-associated events, such as calcium-dependent phospholipid binding, calcium ion binding, negative regulation of calcium ion transport, calcium-dependent protein kinase regulator activity, and calcium-dependent activation of synaptic vesicle fusion (Figure 7A). We thus explored how SELENOT modulates the Ca^2+^ signaling. *SELENOT* siRNA knockdown reduced steady-state cytosolic Ca^2+^ levels in HEK293 cells (Figure 7, B-C). Three Ca^2+^ channels are known to locate on the ER membrane, i.e., sarco-ER Ca^2+^ ATPase (SERCA), ryanodine receptor (RYR) and inositol 1,4,5 trisphosphate receptor (IP3R); among them, SERCA mediates Ca^2+^ influx to ER whereas RYR and IP3R mediate Ca^2+^ efflux to cytosol (14). Next, three available chemical interferers of Ca^2+^ channels [cyclopiazonic acid (CPA), a SERCA antagonist; caffeine, a RYR agonist; adenophostin A (AdA), an IP3R agonist) were employed. *SELENOT* siRNA knockdown in HEK293 cells reduced the peak increment of cytosolic Ca^2+^ concentration when treated with CPA (15 µM; Figure 7D) but not caffeine or AdA treatment (10 mM and 1 µM, respectively; Figure 7, E and F), Conversely, *SELENOT* overexpression increased steady-state cytosolic Ca^2+^ levels (Figure 7, G and H) and enhanced the peak increment in response to CPA treatment in the cells (15 µM; Figure 7I). Likewise, results of Ca^2+^ imaging using AAV-Gcamp 6s-infected brain slices showed a reduction in steady-state cytosolic Ca^2+^ levels (Figure 7, J and K) and a suppression of the peak increment in response to CPA treatment (30 µM; Figure 7L) in the *Selenot^fl/fl^;Dat-cre* slices compared to *Dat-cre* controls. Further analyses in HEK293 cells demonstrated that CPA treatment indeed time- and dose-dependently upregulated the expression levels of *NURR1* and *DAT* (Figure 7, M and N). Together, these results suggest that SELENOT increases cytosolic Ca^2+^ levels via inhibiting SERCA activity, and the increase of cytosolic Ca^2+^ promotes *NURR1* and *DAT* expression.

**Figure 7.**
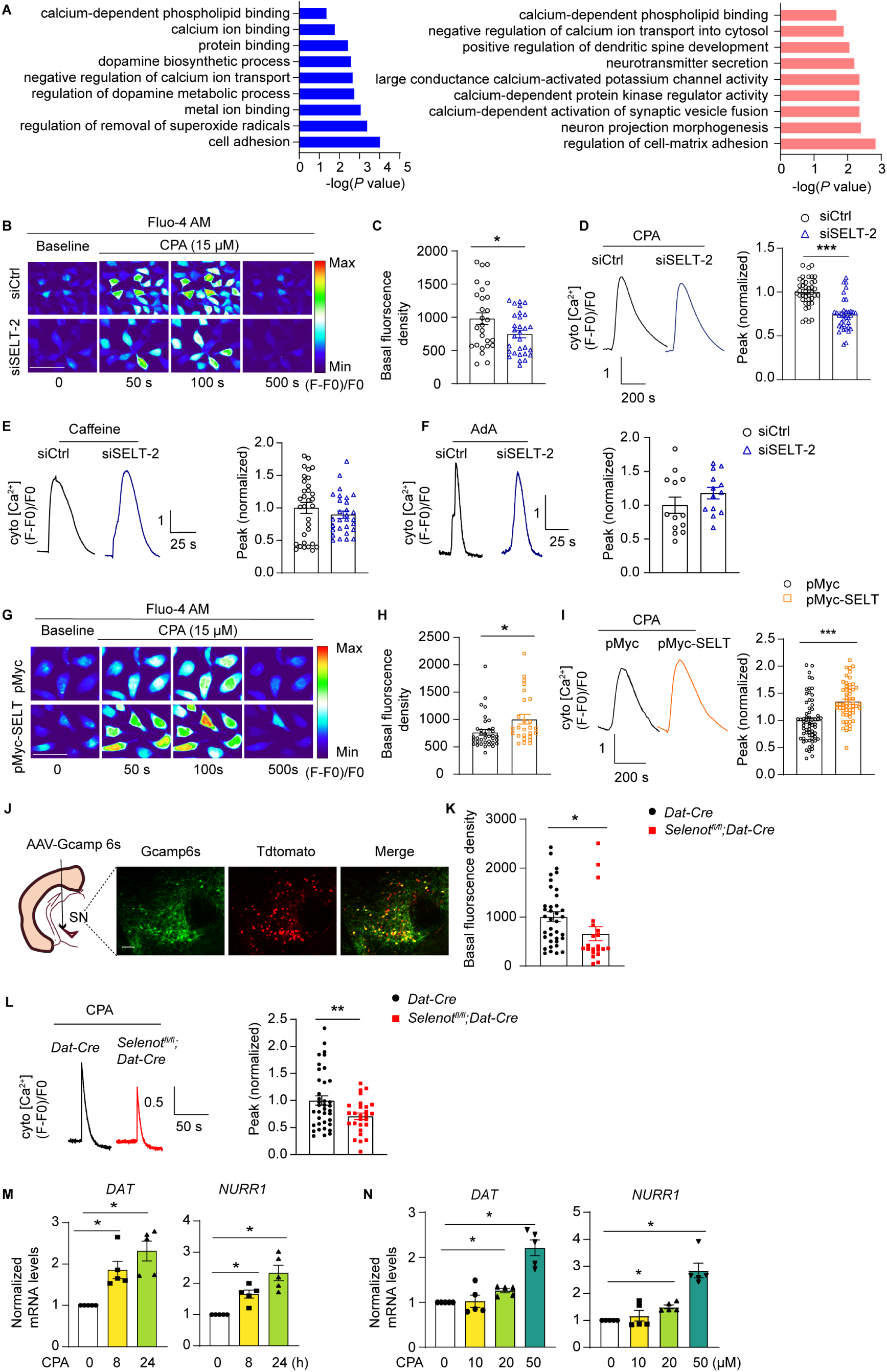
SELENOT modulates ER-cytosol Ca^2+^ influx via SERCA channel. (A) Gene Ontology enrichment analysis for the DEGs. Blue, upregulated; Pink, downregulated; *P* < 0.05. **(B**-**F)** Cytosolic Ca^2+^ level analyses of HEK293 cells transfected with siCtrl or siSELT- 2 for 48 h. Presented are representative images (**B**), steady-state levels (**C**), CPA-induced peaks (**D**), caffeine-induced peaks (**E**), and AdA-induced peaks (**F**). Data were quantified from 3 independent experiments, n = 27 siCtrl and 30 siSELT-2 cells for (**C**), n = 30 siCtrl and 29 siSELT-2 cells for (**D**), n = 32 siCtrl cells and n = 29 siSELT-2 cells for (**E**), n = 13 siCtrl cells and n = 13 siSELT-2 cells for (**F**). Scale bar, 50 μm. **(G**-**I)** Cytosolic Ca^2+^ level analyses of HEK293 cells transfected with pSELENOT or EV for 48 h. Presented are representative images (**G**), steady-state levels (**H**), and CPA-induced peaks (**I**). Data were quantified from 3 independent experiments, n = 36 pMyc and 26 pMyc-SELT cells for (**H**), n = 65 pMyc and 58 pMyc-SELT cells for (**I**). Scale bar, 50 μm. **(J**-**L)** Cytosolic Ca^2+^ level analyses of the substantia nigra slices of mice injected with cre-initiated AAV-Gcamp 6s. Presented are injection diagram and representative images (**J**), steady-state levels (**K**), and CPA-induced peaks (**L**). Data were quantified from 3 *Dat-cre* and 3 *Selenot^fl/fl^;Dat-cre* mice, n = 37 *Dat-cre* and 21 *Selenot^fl/fl^;Dat-cre* cells for (**K**), n = 38 *Dat-cre* and 27 *Selenot^fl/fl^;Dat- cre* cells for (**L**). Red (Tdtomato) positions nuclei. Scale bar, 100 μm. (**M-N**) Time- (**M**) and dose-dependent (**N**) effects of CPA on *DAT* and *NURR1* expression. HEK293 cells were treated with 50 μM CPA for 0, 8 and 24 h (**M**), or with 0, 10, 20, and 50 μm CPA for 24 h (**N**). Quantifications are normalized to β-actin. n = 5 independent experiments. Data are presented as means ± SEM and analyzed by two-tailed unpaired *t*-test for (**B**-**L**), and one-way ANOVA followed by Dunnett’s post-hoc test for (**M** and **N**). \**P* < 0.05; \*\**P* < 0.01; \*\*\**P* < 0.001. AAV, adeno-associated virus; AdA, adenophostin A; CPA, cyclopiazonic acid; DEG, differentially expressed gene; pMyc, pCMV-Myc empty vector; pMyc-SELT, Myc-tagged SELENOT vector; siCtrl, scramble siRNA; siSELT, *SELENOT* siRNA.

### SELENOT physically interacts and colocalizes with SERCA2 in the ER

We thus performed co-immunoprecipitation coupled with mass spectrometry to identify potential SELENOT-interacting proteins, wherein SELENOT^U49C^ mutant was used to achieve higher expression amount as bait protein (Figure 8A). The assay identified 242 unique potential SELENOT^U49C^-interacting proteins with protein scores over 30. Among the isoforms of three ER channels, SERCA, IP3R and RYR, we found that SERCA2 was significantly enriched with 41 peptides and protein score at 188, but none else being qualified (Figure 8, B and C). Interestingly, *Serca2*/*Atp2a2*, but not *Serca1*/*Atp2a1* and *Serca3*/*Atp2a3*, was abundantly expressed in the substantia nigra as suggested by the RNA-seq results. In addition, *Ip3r1* and *Ryr2* were expressed relatively higher than their fellow isoforms in this nucleus (Figure 8D). We measured the expression levels of SERCA2, IP3R1 and RYR2 using *Selenot^fl/fl^;Nestin-cre* and *Selenot^fl/fl^* cortices and found that *Selenot* deficiency did not lead to alteration in their expression (Supplementary Figure 7, B and C). In addition, the SELENOT- interacting proteins were found to be enriched in a number of human phenotypes and diseases, including hyperactivity, neurodevelopmental abnormality, and central nervous system disease (Figure 8E).

**Figure 8.**
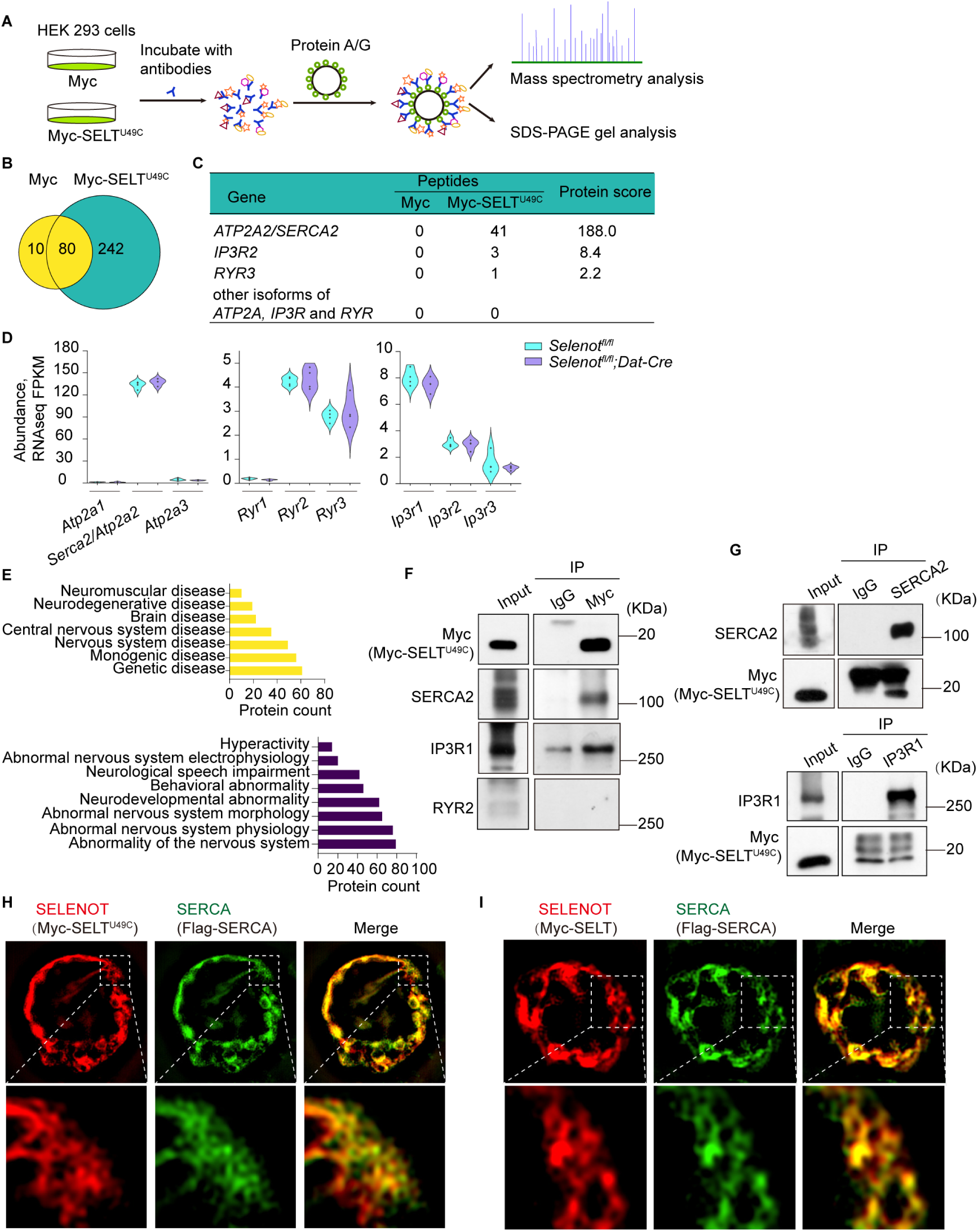
SELENOT interacts with SERCA2, but not IP3R1 and RYR2. (**A**) Schematic diagram to identify SELENOT-interacting proteins. (**B**) Venn diagram of proteins identified by mass spectrometry of immunoprecipitants from Myc and Myc-SELT^U49C^. Numbers refer to number of proteins pulled down with each bait protein (Protein score > 30). (**C**) Enriched peptide numbers and protein scores of ATP2A/SERCA, IP3R, and RYR isoforms. (**D**) Violin plot of the substantia nigra RNA-seq FPKMs of *Atp2a*/*Serca*, *Ip3r*, and *Ryr* isoforms. (**E**) Clustering of enriched human phenotypes and diseases among the 242 unique SELENOT- interacting proteins. (**F**) IP with Myc-SELT^U49C^ to detect SERCA2, IP3R1 and RYR. (**G**) IP with SERCA2 or IP3R1 to detect Myc-SELT^U49C^. HEK293 cells were transfected with pMyc-SELT^U49C^ or pMyC for 48 h. n = 3 independent experiments. (**H** and **I**) Colocalization analyses of SERCA2 with Myc-SELT^U49C^ (**H**) and Myc-SELT (**I**). HEK293 cells were co- transfected with plasmids pFlag-SERCA and pMyc-SELT^U49C^ or pMyc-SELT for 48 h. n = 3 independent experiments. flag-SERCA, flag-tagged SERCA; FPKM, fragments per kilobase per million; IP, immunoprecipitation; IP3R, inositol 1,4,5-triphosphate receptor; pFlag- SERCA2, Flag-tagged SERCA2 vector; pMyc, pCMV-Myc empty vector; pMyc-SELT, Myc- tagged SELENOT vector; pMyc-SELT^U49C^, Myc-tagged SELENOT^U49C^ vector; RYR, ryanodine receptor; SERCA, sarco-ER Ca^2+^ ATPase.

We next performed independent co-immunoprecipitation validation using SELENOT^U49C^ as the bait. Results showed that SELENOT^U49C^ could indeed pull down SERCA2, and also appeared to have some interaction with IP3R1, but not RYR2 (Figure 8F). However, reverse co-immunoprecipitation assay showed that SELENOT^U49C^ could be pulled down by SERCA2, but not by IP3R1 (Figure 8G). Further immunofluorescence colocalization analyses confirmed that SERCA2 was colocalized with SELENOT^U49C^ and also with wild-type SELENOT in the ER (Figure 8, H and I).

### Amphetamine and methylphenidate alleviate hyperactivity in *Selenot^fl/fl^;Dat-cre* mice

Two psychostimulants commonly used for ADHD medication, amphetamine and methylphenidate (15), were used to investigate their efficacy to alleviate the *Selenot* deficiency-induced ADHD-like symptoms. Locomotion in the *Selenot^fl/fl^*control mice was sensitized by treatment with methylphenidate (15 and 30 mg/kg; Figure 9, E-H) but not amphetamine (1 or 2 mg/kg; Figure 9, A-D); these results were similar to the reported effects of the psychostimulants on wild-type mice with these dosages (16). Treatment with amphetamine at 2 mg/kg (Figure 9, C and D) or methylphenidate at 30 mg/kg (Figure 9, G and H), but not at lower dosages (Figure 9, A, B, E, and F), rescued the hyperlocomotion phenotype in *Selenot^fl/fl^;Dat-cre* mice. Such results with pharmacological implications further corroborate the novel role of SELENOT in the protection against ADHD.

**Figure 9.**
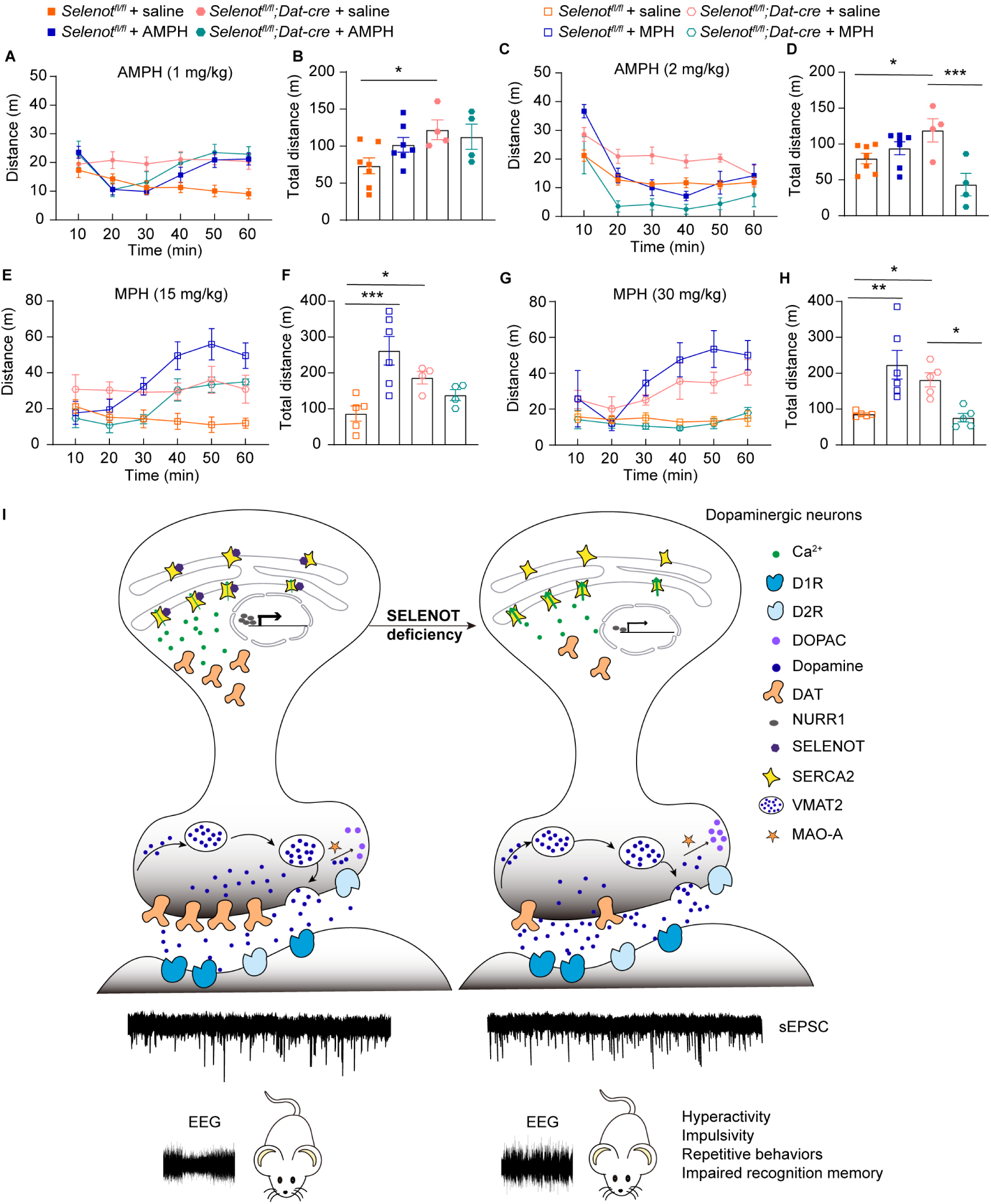
AMPH and MPH ameliorates hyperactivity in *Selenot^fl/fl^;Dat-cre* mice. (**A** and **B**) Open field test of mice treated with saline or 1 mg/kg AMPH. n = 7 *Selenot^fl/fl^* + saline mice, n = 7 *Selenot^fl/fl^* + AMPH mice, n = 4 *Selenot^fl/fl^;Dat-cre* + saline mice, and n = 4 *Selenot^fl/fl^;Dat-cre* + AMPH mice. (**C** and **D**) Open field test of mice treated with saline or 2 mg/kg AMPH. n = 7 *Selenot^fl/fl^* + saline mice, n = 7 *Selenot^fl/fl^* + AMPH mice, n = 4 *Selenot^fl/fl^;Dat-cre* + saline mice, and n = 4 *Selenot^fl/fl^;Dat-cre* + AMPH mice. (**E** and **F**) Open field test of mice treated with saline or 15 mg/kg MPH. n = 5 *Selenot^fl/fl^* + saline mice, n = 6 *Selenot^fl/fl^* + MPH mice, n = 4 *Selenot^fl/fl^;Dat-cre* + saline mice, and n = 4 *Selenot^fl/fl^;Dat-cre* + MPH mice. (**G** and **H**) Open field test of mice treated with saline or 30 mg/kg MPH. n = 5 *Selenot^fl/fl^* + saline mice, n = 6 *Selenot^fl/fl^* + MPH mice, n = 5 *Selenot^fl/fl^;Dat-cre* + saline mice, and n = 5 *Selenot^fl/fl^;Dat-cre* + MPH mice. Presented are locomotor activity every 10 min (**A**, **C**, **E**, and **G**) and total distance for 60 min (**B**, **D**, **F**, and **H**). Data are presented as means ± SEM and analyzed by factorial ANOVA. \**P* < 0.05; \*\**P* < 0.01; \*\*\**P* < 0.001. (**I**) Working model for SELENOT deficiency inducing ADHD behaviors. AMPH, amphetamine; DAT, dopamine transporter; DOPAC, 3,4-dihydroxyphenylacetic acid; D1R, dopamine D1 receptor; D2R, dopamine D2 receptor; EEG, electroencephalogram; MAO-A, monoamine oxidase; MPH, methylphenidate; NURR1, nuclear receptor-related 1; SELENOT, selenoprotein T; SERCA2, sarco-ER Ca^2+^ ATPase 2; sEPSC, spontaneous excitatory postsynaptic currents; VMAT2, vesicular monoamine transporter 2.

## Discussion

ADHD is a prevalent neurodevelopmental disorder affecting millions of children and adults worldwide. Significant progresses have been made into understanding ADHD in etiology, lesioned processes and systems, and treatments. However, a number of uncertainties remains such as to address heterogeneity for precision treatment, to develop new accepted working models, and to identify specific targets for long-term efficacious interventions (11). In the present study, we demonstrate that *Selenot* deficiency in dopaminergic neurons leads to ADHD behaviors in mice. The hyperactivity is ameliorable by psychostimulant treatment.

The underlying pivotal mechanisms towards ADHD are the release of SELENOT-inhibited SERCA pump activity on the ER membrane and the subsequent reduced cytosolic Ca^2+^ levels, which sequentially ablate NURR1 and DAT expression; the DAT defect together with MAO- A impairs presynaptic dopamine reuptake and elevates dopamine metabolism, leading to enhanced extrasynaptic dopamine retention and increased postsynaptic neuronal excitability in the striatum (Figure 9I).

A few rodent models of ADHD targeting different genes have been reported, and they carry differential ADHD-relevant symptoms (12). For instance, high impulsive rats show impulsive behavior and deficits in premature responding with reduced DRD2 expression in nucleus accumbens (17, 18). G protein-coupled receptor kinase interacting protein-1 (*GIT1*) knockout mice display hyperactivity and impaired learning and memory with enhanced cortical EEG theta rhythms, and reduced brain RAC1 signaling and hippocampal inhibitory presynaptic input (19). The earliest ADHD model presumably is made from the DAT knockout mice, which exhibit hyperactivity and impaired cued and spatial learning in association with persistent extracellular hyperdopaminergic tone (20–22). In the present study, phenotypes of *Selenot^fl/fl^;Dat-cre* mice include hyperactivity, recognition memory deficit, repetitive movement, and impulsivity, and they are sensitive to amphetamine or methylphenidate treatment. Nonetheless, no marked impairment is found in anxiety, spatial memory, motor coordination, or social interaction. Phenotypes of *Selenot^fl/fl^;Nestin-cre* mice mostly resemble those of *Selenot^fl/fl^;Dat-cre* mice, in particular the hyperactivity. In contrast, *Selenot^fl/fl^;Gfap-cre* mice carry no ADHD-like behaviors such as hyperactivity and repetitive movement. These results suggest that the modulation of ADHD behaviors is mainly attributed to SELENOT functions in dopaminergic neurons but not astrocytes. Indeed, the dopaminergic system is one of the key neurotransmission circuits disturbed in ADHD, in particular as manifested by the first-line drugs, AMPH and MPH, both of which target this system (23).

Results from previous models of ADHD suggest pathological defects in the monoaminergic system, synaptic transmission, cell adhesion, and signal transduction (12). In the present study, it appears that Ca^2+^ is the direct target of SELENOT, and DAT principally links SELENOT to ADHD. Indeed, both *Selenot^fl/fl^;Dat-cre* mice and *Selenot^fl/fl^;Nestin-cre* mice exhibit drastic reduction in DAT abundance in the VTA, substantia nigra, and striatum, which may lead to the behavioral phenotypes. However, different from *Selenot^fl/fl^;Dat-cre* mice that grow normally, *Selenot^fl/fl^;Nestin-cre* mice show growth retardation as evidenced by reduced body weight and brain size (24), further suggesting a role of SELENOT in the maintenance of brain development. In addition, we speculate that whole-brain *Selenot* deficiency leads to neurocircuit compensation in the brain, which may explain why the ADHD behaviors are less severe in *Selenot^fl/fl^;Nestin-cre* mice than in *Selenot^fl/fl^;Dat-cre* mice. The observation of SELENOT regulation on DAT is corroborated by evidence from cells with SELENOT overexpression and knockdown. We further disclose SELENOT as a positive regulator of NURR1. This nuclear receptor not only transcriptionally activates DAT expression (13), but also is involved in several brain disorders such as Parkinson’s disease (25). Noteworthy, NURR1 deficiency is reported to associate with ADHD-like phenotypes in mice, including hyperlocomotion and impulsivity, but not anxiety and alterations in motor coordination, sociability, and memory (26). As noted earlier, SELENOT is implicated in the modulation of Ca^2+^ flow between ER and cytosol (5, 8). We herein elucidate that the SELENOT-mediated ER-cytosol Ca^2+^ flow in dopaminergic neurons is via the interaction with SERCA2 and then inhibition of its pump activity, but not through the efflux channels, RYRs and IP3Rs. The impact of Ca^2+^ on DAT abundance was less known previously. There was only one study, consistent with our finding, showing that DAT function as determined by dopamine uptake is reduced upon treatment of nifedipine (a L-type Ca^2+^ channel blocker) in mesencephalic cultures (27). In contrast, the transcription factor NURR1 has been noted to be regulatable by nifedipine and L-type calcium channel agonist (28, 29), confirming with us that NURR1 is a target of Ca^2+^ signaling.

According to studies of twins, families, and adoptive siblings, ADHD is highly heritable, ranging from 60% to 90% (12). ADHD patients exhibit EEG abnormalities with most consistent increase in low-frequency activity, predominated by high levels of theta (30–32).

Theta power heritability is the highest in occipital regions in a multivariate analysis of EEG rhythms (33). We herein demonstrate a predominant elevation in occipital theta power, but not in other frequency bands, in *Selenot^fl/fl^;Dat-cre* mice. Nonetheless, no genetically mutated genes have been identified in reported cases of ADHD in humans. Known candidate genes such as *DAT* and *GIT1* are suggested by GWAS, meta-analyses, large-scale linkage studies, and animal model studies (12). *SELENOT* has not yet been documented with pathogenic mutations in human diseases (34). Unfortunately, no brain dataset is available in NCBI GEO to verify the *SELENOT* expression change in human ADHD patients. Results of searching in GWAS datasets indicate a weak association between *SELENOT* variants (such as rs143261451, *P* = 0.014) and ADHD (35). Further verification is needed with the use of human brain samples when conditions allow.

Our findings not only suggest an ADHD-causative gene, but also present a novel ADHD animal model with Ca^2+^- and dopamine-centered explorations. DAT knockout mice bear impaired growth and reduced survival rate with premature death (20), which however were not observed in *Selenot^fl/fl^;Dat-cre* mice. Among the behavioral phenotypes, hyperactivity and impulsivity are characterized in both *Selenot^fl/fl^;Dat-cre* mice and DAT knockout mice (21, 36), but they differ in spatial learning and recognition memory. Therefore, the DAT reduction, the diminished dopamine reuptake in presynaptic dopaminergic terminals, and the enhanced postsynaptic neural excitability appear to be the key, but not all, pathological events leading to the phenotype of dopaminergic *Selenot* deficiency. The *Selenot* deficiency causes no damage in presynaptic dopaminergic neural firing, nor in apoptotic and ER cellular stress. The observation of ER stress not being affected is different from that in corticotrope cells where *Selenot* deficiency is known to induce ER stress (10). Likely, a compensatory mechanism is involved in vivo to maintain the ER homeostasis. As noted earlier, the slight decrease of dopaminergic neurons in the midbrain is likely resulted from proliferation arrest of progenitors during embryonic development (8), but such amount of decrease does not impact the nigrostriatal innervation and dopamine synthesis. Another important finding from the current study is the key role of Ca^2+^ homeostasis in ADHD. Such a role has not been previously underlined, except occasional genetic association reports between ADHD and Ca^2+^-related genes such as those encoding voltage-gated calcium channels (37, 38), as well as a report showing lower serum Ca^2+^ level in ADHD children than normal ones (39).

Additional changes in the dopaminergic system include the MAO-A upregulation and the enhanced dopamine metabolism. As a note, dopamine is metabolized primarily by MAO- B in humans but by MAO-A in rodents (40–42). It remains unclear as to what the increased dopamine metabolism means to ADHD, but brain MAO-A elevation is associated with a variety of psychiatric illnesses and prodromal states as suggested by brain imaging and post-mortem evidence (43), and MAO-A polymorphisms are known to be associated with ADHD susceptibility in human populations (44).

In conclusion, midbrain SELENOT is indispensable for maintenance of proper dopamine signaling. SELENOT deficiency in dopaminergic neurons, but not in astrocytes, leads to ADHD in mice, the pathology of which is mechanistically associated with a cascade involving the lost interaction and inhibition of SERCA, the reduced cytosolic Ca^2+^ levels, the ablation of NURR1 and DAT, the reduced dopamine reuptake and enhanced extra synaptic dopamine retention, and the enhanced striatal neurotransmission. Our findings suggest that SELENOT has a protective role against ADHD pathogenesis and may be a target for patient genetic screening and therapeutics.

## Materials and Methods

### Mice and chemicals

The C57BL/6 line of *Selenot^fl/fl^* mice was generated by flanking *Selenot* exon 2 and exon 3 with *loxP* sites (Cyagen Inc., Guangzhou, China). The *Dat/Slc6a3-cre, Nestin-cre*, and *Gfap-cre* mice (Cyagen) contain Cre recombinase knock-in alleles driven by specific promoter for gene knockout in dopaminergic neurons, in the central and peripheral nervous system including neuronal and glial cell precursors, and in astrocytes, respectively; they were cross- bred with *Selenot^fl/fl^* mice to generate *Selenot^fl/fl^;Dat-cre, Selenot^fl/fl^;Nestin-cre*, and *Selenot^fl/fl^;Gfap-cre* mice. Mice (3-4 per cage) were given free access to pelleted chow diet and distilled water in a specific pathogen-free animal facility maintained at 22℃ and 40-60% humidity and with a 12-h light-dark cycle. Unless otherwise indicated, male mice aged 8-12 weeks were used and sacrificed under deep isoflurane anesthesia. All the mouse experiments were approved by the Institutional Laboratory Animal Care and Use Committee of Wenzhou Medical University and conducted in accordance with China National Standard Guidelines for Laboratory Animal Welfare. PCR genotyping for *cre* and *Selenot* was performed using primer pairs listed in Supplementary Table 1. AMPH (A-007), caffeine (C0750), dopamine (H8502), nimodipine (N149) and CPA (C1530) were purchased from Sigma, MPH (298-59-9) from Rhawn Reagent, AdA (sc-221213) from Santa Cruz Biotechnology, and Biocytin (28022) from Thermo Fisher Scientific.

### Behavioral assessments

Behavior parameters were recorded and analyzed using the ANY-maze video tracking system (Stoelting Co.). For open-field test, the spontaneous locomotor activity was recorded for 60 min in a 50×50×50 cm open field chamber. For elevated plus maze, the apparatus was opaque and comprised of a central platform (10×10 cm), two open arms (50×10 cm), two closed arms (50×10 cm) with protective walls 40 cm high, and a supporting rod (height, 50 cm). The mouse was placed on an open arm facing another open arm, and the activity was recorded for 5 min. For Y-maze test, the mouse was placed in a Y-shaped maze with three white, opaque arms (35×6 cm) at 120° angle from each other. The mouse was allowed to freely explore the three arms and recorded for 10 min. For rotarod test, the mouse was placed on an open rotarod facing away from the experimenter and trained under the accelerating condition (4 to 40 rpm over 5 min) for three consecutive days. The latency to fall was recorded on day 4.

For three-chamber social analysis, the apparatus was a 60×42×22 cm box divided into 3 equal-sized compartments with doors in between. The mouse was tested in the three chambers for three sessions. At session 1, the test mouse was allowed to freely explore in the center chamber for 5 min to habituate, with doors to the side chambers closed. At session 2, a social mouse (stranger 1) in a wire cage was placed into one of the side chambers and the test mouse was allowed to explore for 10 min in three chambers with doors open. At session 3, a new mouse (stranger 2) in a wire cage was placed into the other side chamber and the test mouse was allowed to freely explore for 10 min. The stimulus mice were age- and sex-matched, and unknown to the test mouse. The time spent on exploring each cage, i.e., within 3-cm range to the cage, was measured.

For novel object recognition, the arena was a 50×50×50 cm open field chamber. On day 1, the mouse was allowed to explore freely in the field for 10 min. On day 2, the mouse was introduced to two identical objects in the field and allowed to explore the objects freely for 10 min. Time of the mouse spending on exploring each object was measured. The mouse was placed back to the field after 1 h (short-term) and 24 h (long-term), with one of the objects replaced with a differently shaped object. Time of the mouse spending on exploring each of the objects was measured. Object exploration was defined when the mouse was within a 3-cm range to the object. Novel object preference is defined as percentage of the novel object exploring time to the total object exploring time. For cliff avoidance reaction, the apparatus was a round, plastic platform (diameter, 20 cm; thickness, 2 cm) stably supported by a plastic rod (height, 50 cm). The mouse was placed on the edge of the platform. Impulsivity was expressed as frequency of the mouse falling from the platform recorded within 60 min.

### Collection of brain tissues

Mice were euthanized and transcardially perfused with 10 mL of saline. The brain was dissected and rinsed with cold water and placed immediately in a chilled rodent brain matrix (RWD Life Science, China). The brain was then sliced with a cold razor blade, starting around 1 mm rostral to the lambda and sectioning with 1 mm thickness for the substantia nigra and striatum. The desired tissue was collected from the slices and fast frozen in liquid nitrogen before stored at -80 ℃ for further use.

### Transfection

HEK293 cells (American Type Culture Collection, Manassas, VA) were cultured in DMEM supplemented with 10% fetal bovine serum (086-150, Wisent) and maintained in a humidified incubator at 37°C with 5% CO_2_. Cells were transfected with siRNA or plasmids for desired time as detailed in the figure legends using Lipofectamine 3000 (L3000015, Invitrogen).

SELENOT siRNAs (Supplementary Table 1) were purchased from GenePharma (Shanghai, China). The Myc-tagged SELENOT expression vector (pMyc-SELT) was constructed with the use of the pCMV-Myc plasmid (pMyc) as previously described (8). The myc-tagged SELENOT^U49C^ vector (pMyc-SELT^U49C^) was made by the method of overlap extension PCR. The flag-tagged SERCA2 vector (pFlag-SERCA2) was purchased from WZ Biosciences (Shandong, China).

### Western blot

Cells or tissues were lysed in a sample buffer (60 mM Tri-HCl, pH 6.8; 2% SDS; 5% glycerol) and boiled for 10 min. After centrifugation, total protein concentration was measured using a BCA kit (P0010, Beyotime). An equal amount of total protein from each sample was loaded and analyzed by Western blot as previously described (45). Primary and secondary antibodies are detailed in Supplementary Table 2. The LumiGLO® Reagent and Peroxide kit (7003) was purchased from Cell Signaling Technology.

### Co-immunoprecipitation

After transfection with pMyc or pMyc-SELT^U49C^ for 48 h, HEK293 cells were lysed in the lysis buffer supplemented with protease and phosphatase inhibitors (P0013, Beyotime). Cell lysates were incubated with IgG or antibodies against Myc, SERCA2, or IP3R1 at 4℃ overnight. An equal volume of protein A/G agarose beads was then added into the lysates and continued for incubation at 4°C for 2 h. After centrifugation, the precipitated beads were washed with PBS for 5 times. The bound proteins were eluted with 1× loading buffer and sent to BGI genomics (Shenzhen, China) to identify proteins with liquid chromograph liquid chromatography tandem mass spectrometry (LC-MS/MS). The expression and immunoprecipitation of Myc-SELENOT were confirmed by both silver staining and Western blot in pre-experiments.

### Real-time PCR

RNA was isolated using RNAiso Plus (9108, Takara) and reversely transcribed to cDNA using the HiScript RT SuperMix (R222-01, Vazyme). Quantitative levels of mRNA were assayed in triplicates using the Universal SYBR Master Mix (MQ101-00, Vazyme) following the manufacturer’s instruction in the CFX Connect System (Bio-Rad Laboratories). Primers for *DAT* and *ACTB* are listed in Supplementary Table 1. Expression levels were calculated as 2^−ΔΔCt^, wherein *ACTB* was an internal control.

### Immunohistochemistry and immunofluorescence

Mice were euthanized and transcardially perfused sequentially with saline (10 mL) and 4% paraformaldehyde (5 mL). Brains were removed and fixed in 4% paraformaldehyde for 24 h before being dehydrated in ethanol, embedded in paraffin, and sectioned to 5 μm. Endogenous peroxidase activity was blocked with 3% H_2_O_2_ at room temperature for 20 min. The brain slices were blocked in 5% bovine serum albumin for 30 min and incubated with primary antibodies at 4°C overnight. For immunohistochemistry, the slices were incubated with horseradish peroxidase-conjugated secondary antibodies for 1 h at room temperature. 3,3’- diaminobenzidine was applied to visualize immunohistochemical staining. Nuclei were counterstained with hematoxylin before mounted with neutral gum. For immunofluorescence, the slices were incubated with secondary antibodies for 2 h at room temperature, and counterstained with Hoechst 33258 for 5 min to locate the nuclei before mounted with anti- fade mounting media. Primary and secondary antibodies are listed in Supplementary Table 2. Images were visualized by Eclipse Ti or A1R-SIM-STORM microscope (Nikon, Japan) and analyzed by ImageJ.

### Calcium imaging

Because the adeno-associated virus (AAV), pAAV-EF1a-DIO-GCaMP6f-P2A-NLS-dTomato (OBiO Technology, Shanghai, China), was initiated for expression under the control of Cre recombinase, *Dat-cre* mice, instead of *Selenot^fl/fl^*mice, were used as a control herein. Mice were stereotaxically injected with 10 μL of AAV (3 × 10^12^ vector genomes/mL) in the substantia nigra [from bregma: anterior-posterior (AP) -3.0 mm, media lateral (ML) +1.2 mm, and dorsal-ventral (DV) -4.5 mm] and sacrificed 3 weeks later. Transverse brain slices were prepared as described in the above electrophysiology section. Slices containing the substantia nigra were recovered in a holding chamber containing oxy-aCSF without CaCl_2_ for 8 min at 33°C. Time lapse images were then recorded at 15 frames per second using the A1R/N-SIM/N-STORM microscope (Nikon), with recording for 1 min at basal levels, followed by drug additions and incubation for 7 min and then recording for 3 min. Images were analyzed by ImageJ.

For cellular calcium imaging, cells were washed and incubated with 2 μM Fluo-4-AM (S1060, Beyotime) in HBSS (137 mM NaCl, 0.3 mM Na_2_HPO_4_·12H_2_O, 5.4 mM KCl, 0.4 mM KH_2_PO_4_, 5.6 mM D-Glucose, 4.2 mM NaHCO_3_) for 30 min at 37°C. After washing with HBSS 3 times, cells were proceeded for time lapse imaging as above described by recording for 1 min at basal levels and 5 min after drug administration.

### Electrophysiology

Transverse brain slices (300 μm thick) were freshly prepared in an ice-cold dissection buffer (211 mM sucrose, 3.3 mM KCl, 1.3 mM NaH_2_PO_4_, 0.5 mM CaCl_2_, 10 mM MgCl_2_, 26 mM NaHCO_3_, and 11 mM glucose) using a vibratome (VT1000S, Leica). Slices were recovered in a holding chamber containing oxygenated (95% O_2_ and 5% CO_2_) artificial cerebrospinal fluid (i.e., oxy-aCSF; 124 mM NaCl, 3.3 mM KCl, 2.5 mM CaCl_2_, 1.5 mM MgCl_2_, 1.3 mM NaH_2_PO_4_, 26 mM NaHCO_3_, and 11 mM glucose) for 1 h at 33 °C. According to the morphology and location, the dopaminergic neurons in the substantia nigra were recorded using borosilicate glass pipettes with resistances in the 5-8 MΩ range, and the pipettes were pulled on a Flaming-Brown micropipette puller (P-1000, Sutter Instruments). Data were acquired and analyzed by pClamp10 and Clampfit 10 in MultiClamp 700B amplifier and 1550 B digitizer, respectively (Molecular Device).

### EEG recording

Mice were anesthetized with 1% (w/v) sodium pentobarbital. Epidural electrodes were bilaterally fixed onto the skull with screws and dental cements. Electromyogram electrodes were placed into neck muscles. After recovery for 1 week, EEG recording was performed on the freely moving mouse at 500 Hz using the EEG/EMG system (Pinnacle). Recorded traces were highpass-filtered with 3 Hz and the first hour was analyzed for power spectral density.

### RNA sequencing

Total RNA was extracted using RNAiso Plus (9108, Takara) based on the manufacturer’s protocol. RNA sequencing was performed by using the Illumina HiSeq X10 system at LC Sciences (Hangzhou, China) as previously described (46). After removing low-quality reads, HISAT package was used to map the reads to the *Mus musculus* reference genome (http://genome.ucsc.edu/). The mapped reads were assembled using StringTie and merged to reconstruct a comprehensive transcriptome using Perl scripts. StringTie was then used to determine mRNA expression levels by calculating fragments per kilobase per million mapped reads. Differentially expressed mRNAs were selected by Ballgown with fold change > 1.2 and *P* < 0.05. Bioinformatic analyses were performed using the OmicStudio tools (https://www.omicstudio.cn/tool) with statistical significance set at *P* < 0.05.

### In vivo microdialysis

The mouse was stereotaxically implanted with a microdialysis probe (CMA-7, Harvard Bioscience) under 1% (w/v) sodium pentobarbital anesthesia in the striatum (from bregma: AP +0.6 mm, ML -1.8 mm, and DV -4.0 mm). After a 12-h recovery, the probe was perfused continuously with artificial cerebrospinal fluid (147 mM NaCl, 3.5 mM KCl, 1.2 mM CaCl_2_, 1.2 mM MgCl_2_, 1 mM NaH_2_PO_4_, and 25 mM NaHCO_3_, pH 7.0-7.4) at a flow rate of 1 μL/min. Ten samples were collected with an interval of 35 min. The first 4 were discarded as equilibration samples, and the other 6 samples were stored at -80°C for dopamine analysis. Location of the probe within the striatum was confirmed histologically.

### Neurotransmitter measurement

Neurotransmitters and metabolites were analyzed by ultra-high-performance LC-MS/MS with quality control assessment. All internal standards were obtained from Sigma. Samples were separated by a UPLC BEH C18 column (100×2.1 mm, 1.7 μm; 186002352, Waters). Striatal sample preparation and measurement were performed at ProfLeader Biosciences (Shanghai, China) using a coupled system of 1290 Infinity II UHPLC and 6470A mass spectrometry (Agilent). Microdialysis samples for dopamine analysis were measured using the QTRAP 6500+ system (AB Sciex).

### Statistical analysis

Statistical differences were evaluated using unpaired two-tailed Student’s *t*-test, Chi square test, one-way analysis of variance (ANOVA) followed by Dunnett’s post-hoc test, repeated measures ANOVA test, or factorial ANOVA test as specified in the figure legends. Data were expressed as mean ± standard error of mean (SEM) from at least three independent experiments. Differences were considered statistically significant at *P* < 0.05. All analyses were conducted with the use of SPSS version 23.0.

## Author contributions

JHZ and XZ conceived the idea; QG charged the project; QG and DYH performed behavioral tests; ZFL, QG, DYH, and KNW investigated mechanisms; PJL and QG performed electrophysiology experiments; HHF aided mouse line generation; HHF, JD, and HMW assisted data analysis and provided resources; JHZ and XZ supervised the study and acquired funding support; JHZ and QG wrote the manuscript; All authors have read, edited, and approved the final manuscript.

## Acknowledgements

The authors are grateful to Dr. Jiang-Fan Chen for assistance in reagents, and the Scientific Research Center of Wenzhou Medical University for consultation and instrument support. This work was supported by Zhejiang Provincial Natural Science Foundation (LD19H090001 and LZ19H090002) and National Natural Science Foundation of China (81771380, 81771510, 82071585, and 82271282).

## Data availability

The bulk RNA-seq data are deposited in the Sequence Read Archive database under the accession number PRJNA910327.

