## Supplementary material for "SELENOT regulates endoplasmic reticulum calcium flux via SERCA2 and maintains dopaminergic DAT to protect against attention deficit hyperactivity disorder in mice": SELENOT manuscript bioRxiv Supplements.docx

**This file includes:**

Supplementary Figures 1 to 7;

Supplementary Tables 1 and 2.

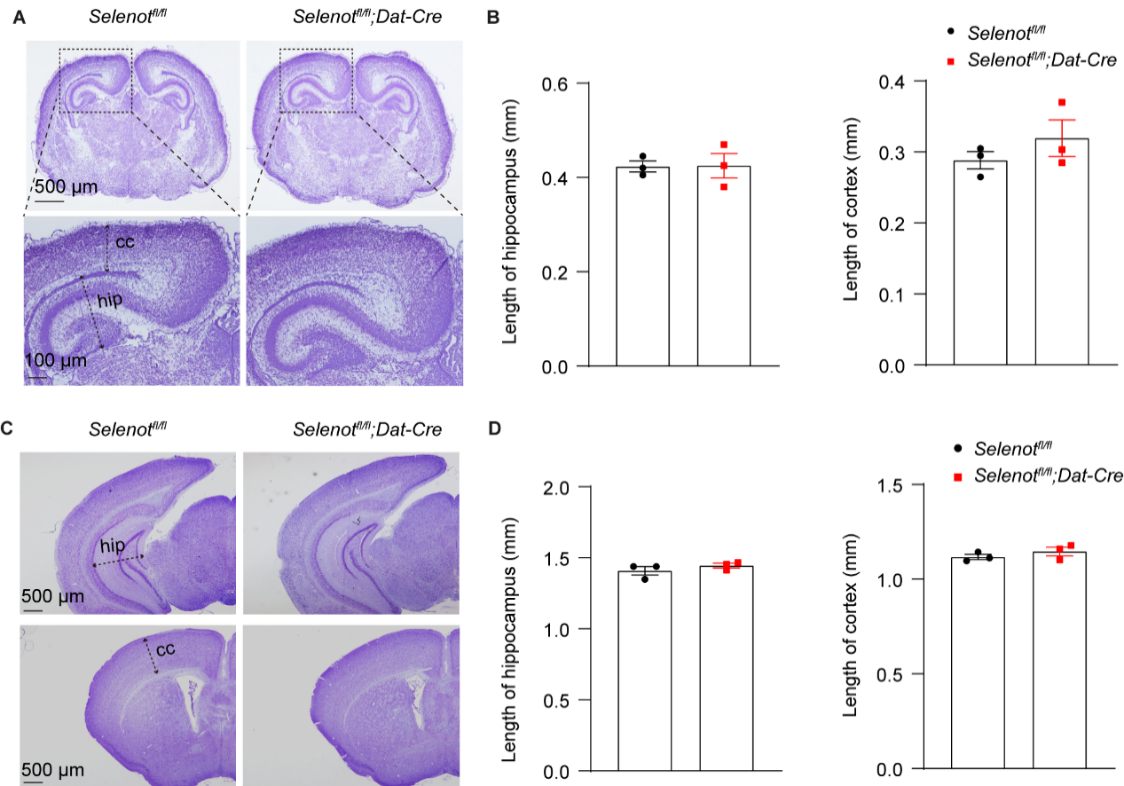

**Supplementary Figure 1. Brain nissl staining.** (**A** and **B**) Representative coronal sections (**A**) and volume quantifications (**B**) of cortex (cc) and hippocampus (hip) in mice aged postnatal 3 days. (**C** and **D**) Representative coronal sections (**C**) and volume quantifications (**D**) of cortex and hippocampus in mice aged 6 weeks. Volume is expressed as the space length indicated by the arrow. Quantification was averaged from 2-4 consecutive slices for each mouse. n = 3 *Selenot^fl/fl^* mice and n = 3 *Selenot^fl/fl^;Dat-cre* mice. Data are presented as means ± SEM and analyzed by two-tailed unpaired *t*-test.

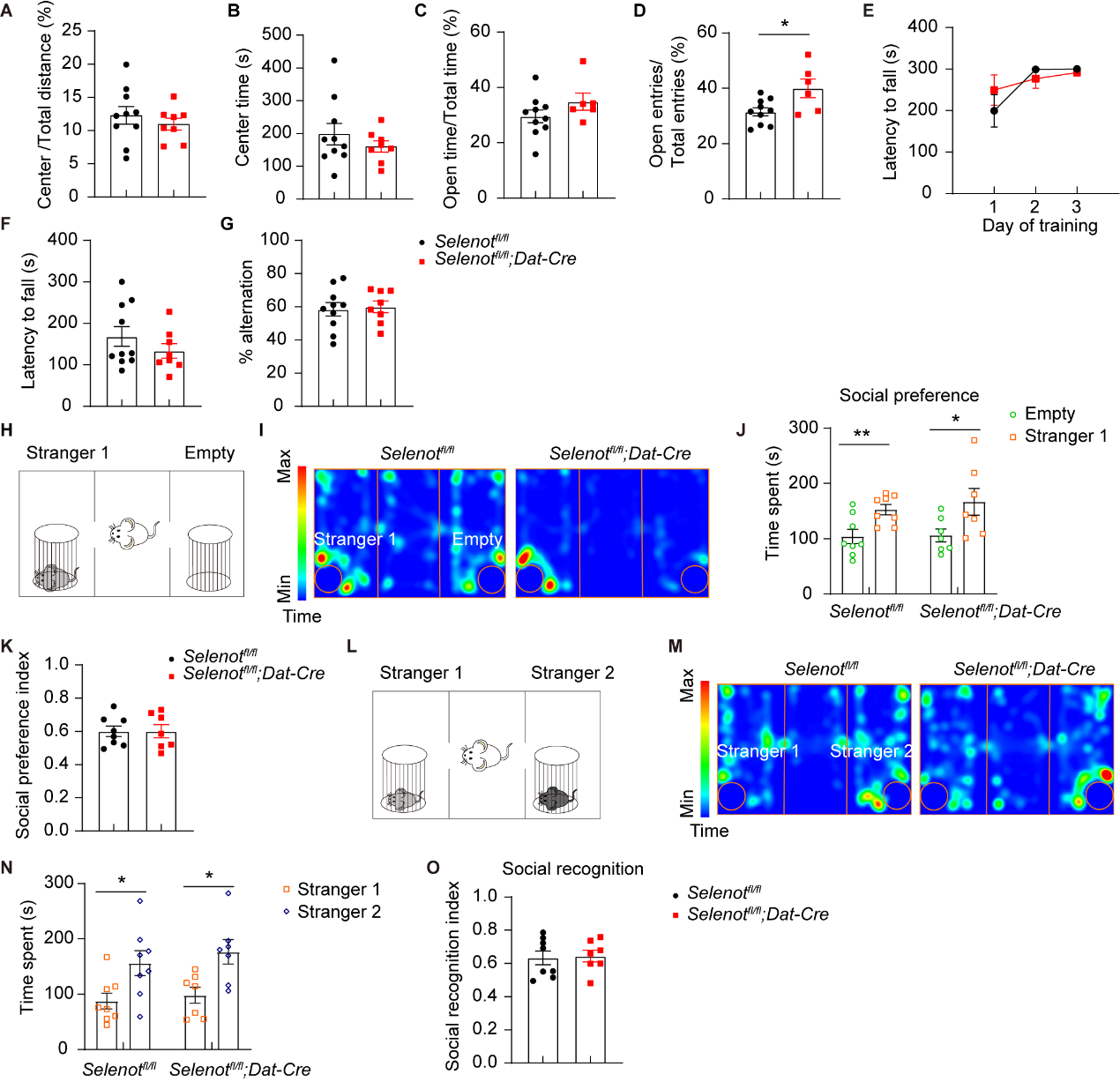

**Supplementary Figure 2. Additional behavioral assessments for *Selenot^fl/fl^;Dat-cre* mice.** (**A** and **B**) Percentage of center distance to total distance (**A**) and center time (**B**) in open field test of *Selenot^fl/fl^* (n = 10) and *Selenot^fl/fl^;Dat-cre* (n = 8) mice. (**C** and **D**) Percentage of time spent in open arms to total time spent in closed and open arms (**C**) and percentage of entries to open arms compared to total entries to open and close arms (**D**) in elevated plus maze test of *Selenot^fl/fl^* (n = 10) and *Selenot^fl/fl^;Dat-cre* (n = 6) mice. (**E** and **F**) Latency to fall in 3-day training (**E**) and test (**F**) in rotarod. n = 10 *Selenot^fl/fl^* mice and n = 8 *Selenot^fl/fl^;Dat-cre* mice. (**G**) Spontaneous alternation in Y maze test. n = 10 *Selenot^fl/fl^* mice and n = 8 *Selenot^fl/fl^;Dat-cre* mice. (**H**-**O**) Three chamber test of *Selenot^fl/fl^* (n = 8) and *Selenot^fl/fl^;Dat-cre* (n = 7) mice. Presented are schematic diagram (**H** and **L**), representative activity heatmap (**I** and **M**), time spent near the cage empty or with a stranger mouse (stranger 1; **J**) and the calculated social preference index (**K**), and time spent near the cage with the familiar mouse (stranger 1) or with another stranger (stranger 2; **N**) and the calculated social recognition index (**O**). Data are presented as means ± SEM and analyzed by two-way repeated measures ANOVA for (**E**) and two-tailed unpaired *t*-test for (**A**-**D** and **F**-**O**). **P* < 0.05; ***P* < 0.01.

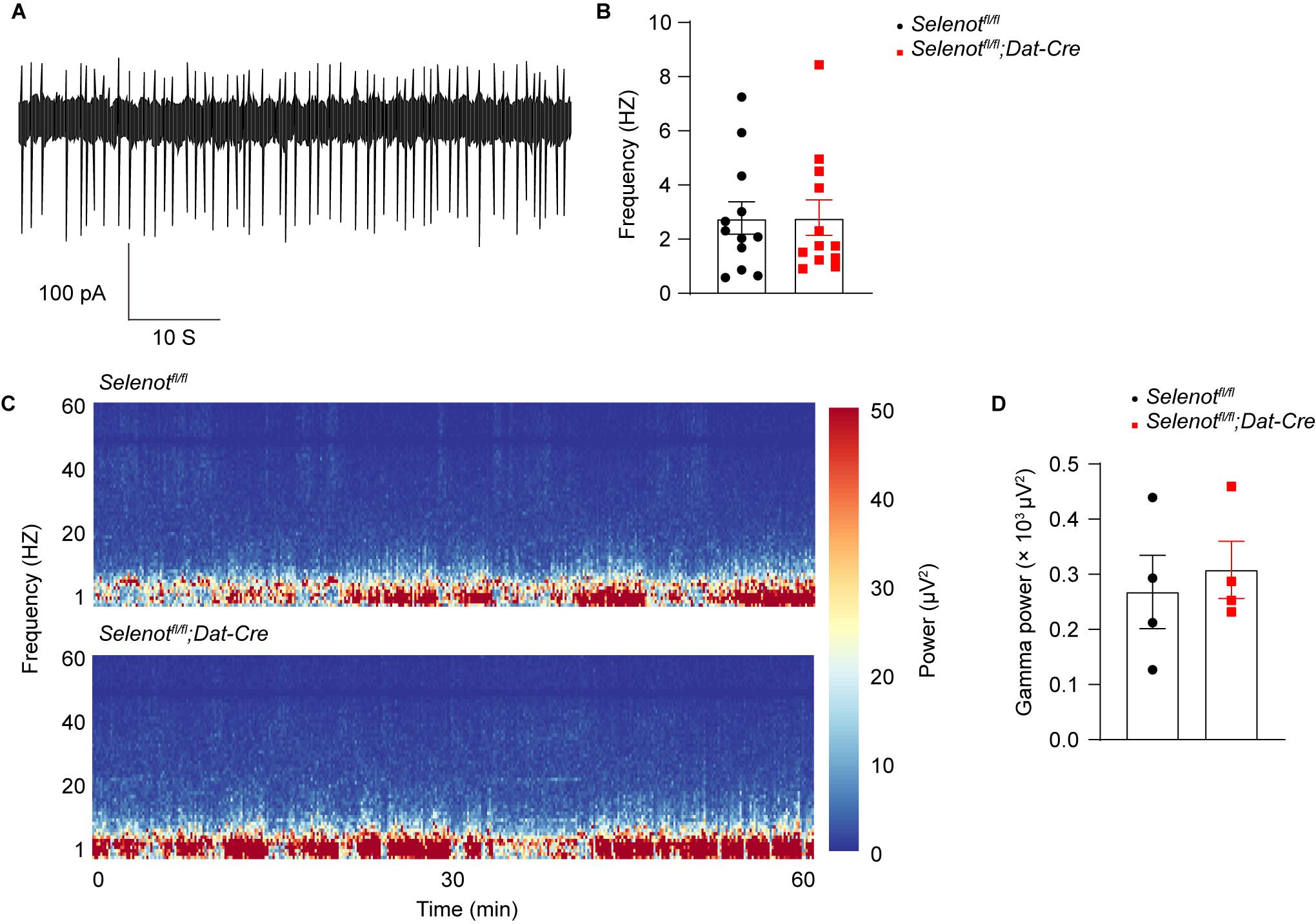

**Supplementary Figure 3. Pacemaker of dopaminergic neurons and EEG power of higher frequency in occipital regions.** (**A** and **B**) Representative traces (**A**) and pacemaker frequencies (**B**) of dopaminergic neurons by cell-attached recording. n = 12 neurons from 3 *Selenot^fl/fl^* mice and n = 12 neurons from 3 *Selenot^fl/fl^;Dat-cre* mice. (**C** and **D**) Representative spectrograms (**C**) and gamma power (27-44 Hz; **D**) in 60 min in occipital regions. n = 4 *Selenot^fl/fl^* mice and n = 4 *Selenot^fl/fl^;Dat-cre* mice. Data are presented as means ± SEM and analyzed by two-tailed unpaired *t*-test. EEG, electroencephalogram.

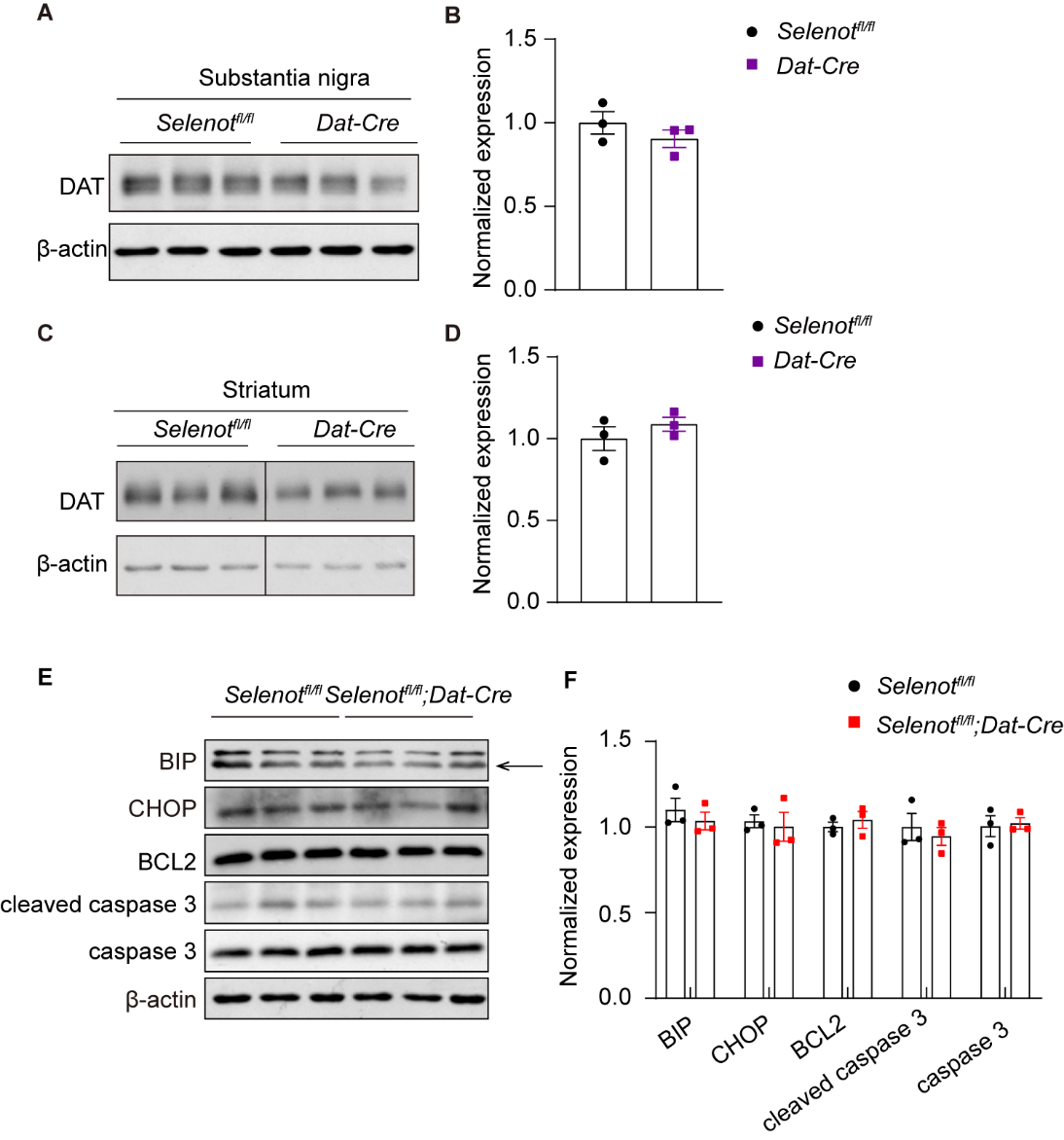

**Supplementary Figure 4. Analyses of DAT expression between *Selenot^fl/fl^* and *Dat-cre* mice and apoptosis and ER stress in *Selenot^fl/fl^;Dat-cre* mice.** (**A-D**) Western blot analyses (**A**, **C**) and quantifications (**B**, **D**) of DAT expression in the substantia nigra and striatum of *Selenot^fl/fl^* mice (n = 3) and *Dat-cre* mice (n =3). Lanes are neighbored from one blot for (**C**). (**E** and **F**) Western blot analyses (**E**) and quantifications (**F**) of proteins associated with apoptosis and ER stress in the substantia nigra of *Selenot^fl/fl^* mice (n = 3) and *Selenot^fl/fl^;Dat-cre* mice (n = 3). Quantifications are normalized to β-actin. Data are presented as means ± SEM and analyzed by two-tailed unpaired *t*-test. BCL-2, B-cell lymphoma-2; BIP, binding-immunoglobulin protein; CHOP, CCAAT-enhancer-binding protein homologous protein; DAT, dopamine transporter; ER, endoplasmic reticulum.

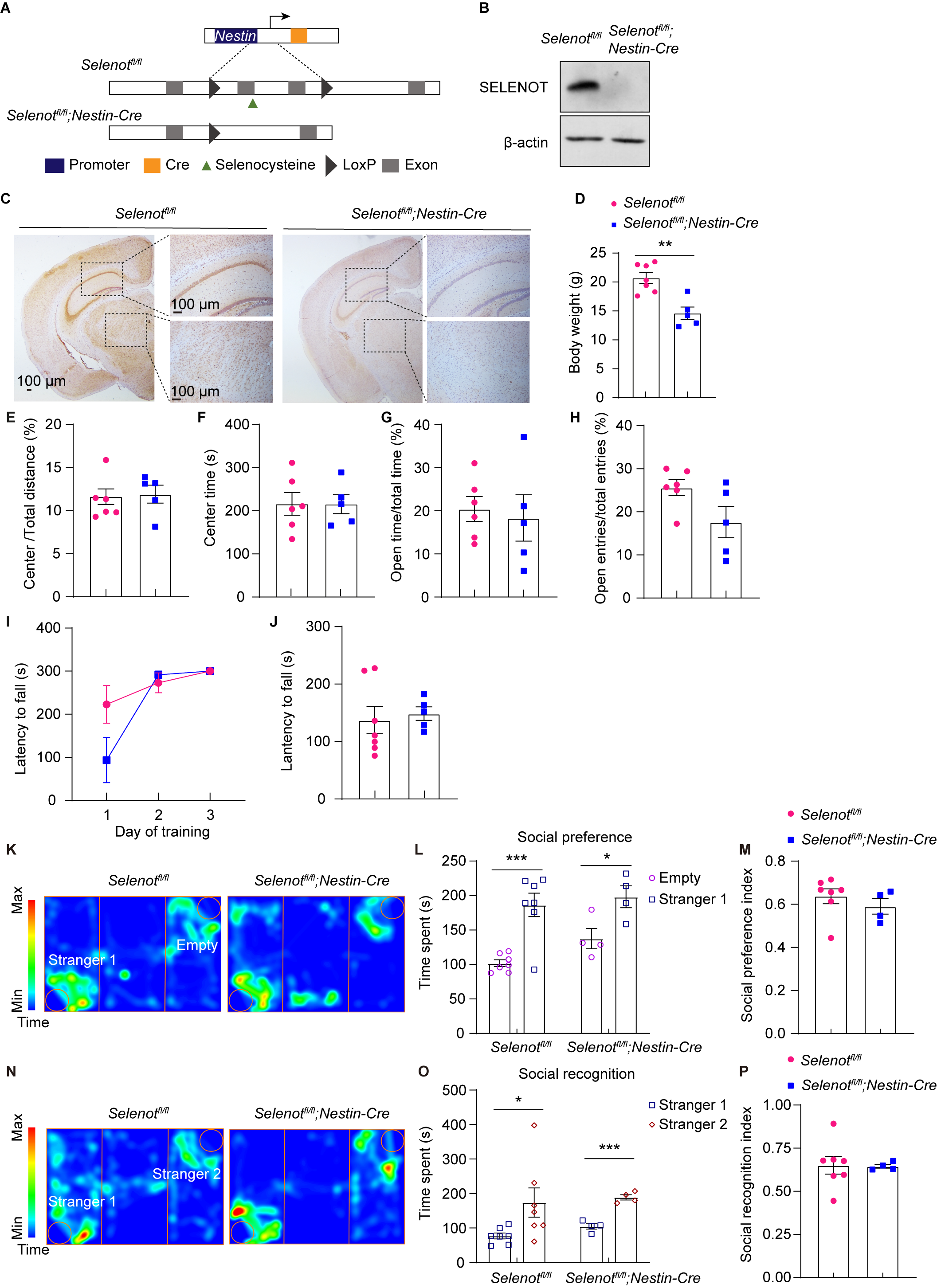

**Supplementary Figure 5. Generation of *Selenot^fl/fl^;Nestin-cre* mice and the additional behavioral assessments.** (**A**) Schematic diagram of *Nestin* promoter-driven excision of *Selenot* exons 2 and 3. (**B** and **C**) Western blot (**B**) and immunohistochemistry (**C**) analyses of SELENOT expression in the brain. (**D**) Body weight of mice at 8 weeks old. n = 7 *Selenot^fl/fl^* mice and n = 5 *Selenot^fl/fl^;Nestin-cre* mice. (**E** and **F**) Percentage of center distance to total distance (**E**) and center time (**F**) in open field test of *Selenot^fl/fl^* (n = 6) and *Selenot^fl/fl^;Nestin-cre* (n = 5) mice. (**G** and **H**) Percentage of time spent in open arms to total time spent in closed and open arms (**G**) and percentage of entries to open arms compared to total entries to open and close arms (**H**) in elevated plus maze test of *Selenot^fl/fl^* (n = 6) and *Selenot^fl/fl^;Nestin-cre* (n = 5) mice. (**I** and **J**) Latency to fall in 3-day training (**I**) and test (**J**) in rotarod. n = 7 *Selenot^fl/fl^* mice and n = 5 *Selenot^fl/fl^;Nestin-cre* mice. (**K**-**P**) Three chamber test of *Selenot^fl/fl^* (n = 7) and *Selenot^fl/fl^;Nestin-cre* (n = 4) mice. Presented are representative activity heatmap (**K** and **N**), time spent near the cage empty or with a stranger mouse (stranger 1; **L**) and the calculated social preference index (**M**), and time spent near the cage with the familiar mouse (stranger 1) or with another stranger (stranger 2; **O**) and the calculated social recognition index (**P**). Data are presented as means ± SEM and analyzed by two-way repeated measures ANOVA for (**I**) and two-tailed unpaired *t*-test for other comparisons. **P* < 0.05; ***P* < 0.01; ****P* < 0.001.

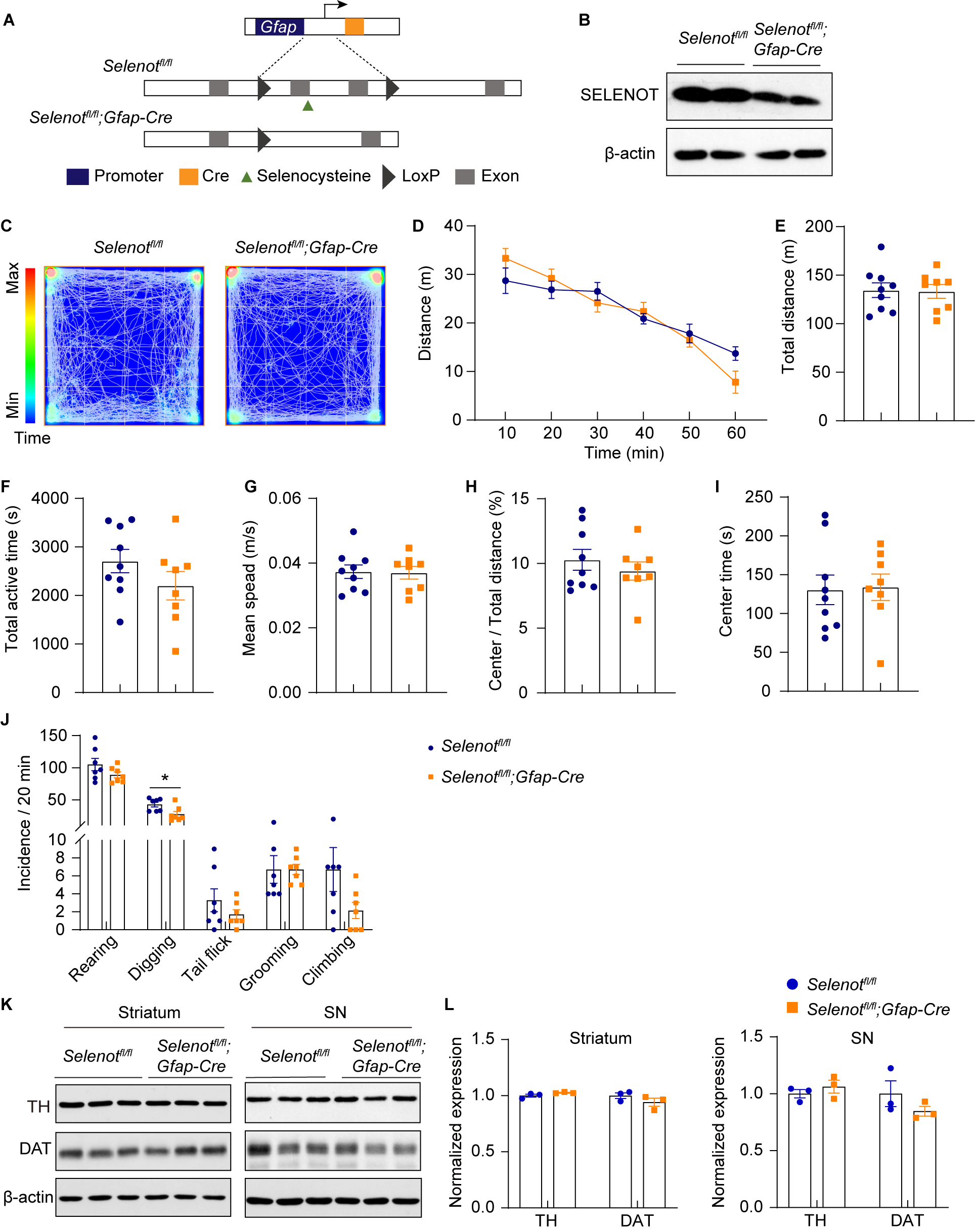

**Supplementary Figure 6. Generation of *Selenot^fl/fl^;Gfap-cre* mice and the locomotion behavior assessments.** (**A**) Schematic diagram of *Gfap* promoter-driven excision of *Selenot* exons 2 and 3. (**B**) Western blot analyses of SELENOT expression using whole brain. (**C**-**I**) Open field test of *Selenot^fl/fl^* (n = 9) and *Selenot^fl/fl^;Gfap-cre* (n = 8) mice. Presented are representative activity heatmap (**C**), distance traveled every 10 min (**D**), total distance traveled (**E**), total active time (**F**), mean speed (**G**), percentage of center distance to total distance (**H**) and center time (**I**). (**J**) Stereotyped behaviors of *Selenot^fl/fl^* (n = 7) and *Selenot^fl/fl^;Gfap-cre* (n = 7) mice. (**K** and **L**) Western blot analyses (**K**) and quantifications (**L**) of TH and DAT expression in the total striatum and substantia nigra. n = 3 *Selenot^fl/fl^* mice and n = 3 *Selenot^fl/fl^;Gfap-cre* mice. Quantifications are normalized to β-actin. Data are presented as means ± SEM and analyzed by two-way repeated measures ANOVA for (**D**) and two-tailed unpaired *t*-test for other comparisons. **P* < 0.05. DAT, dopamine transporter; TH, tyrosine hydroxylase.

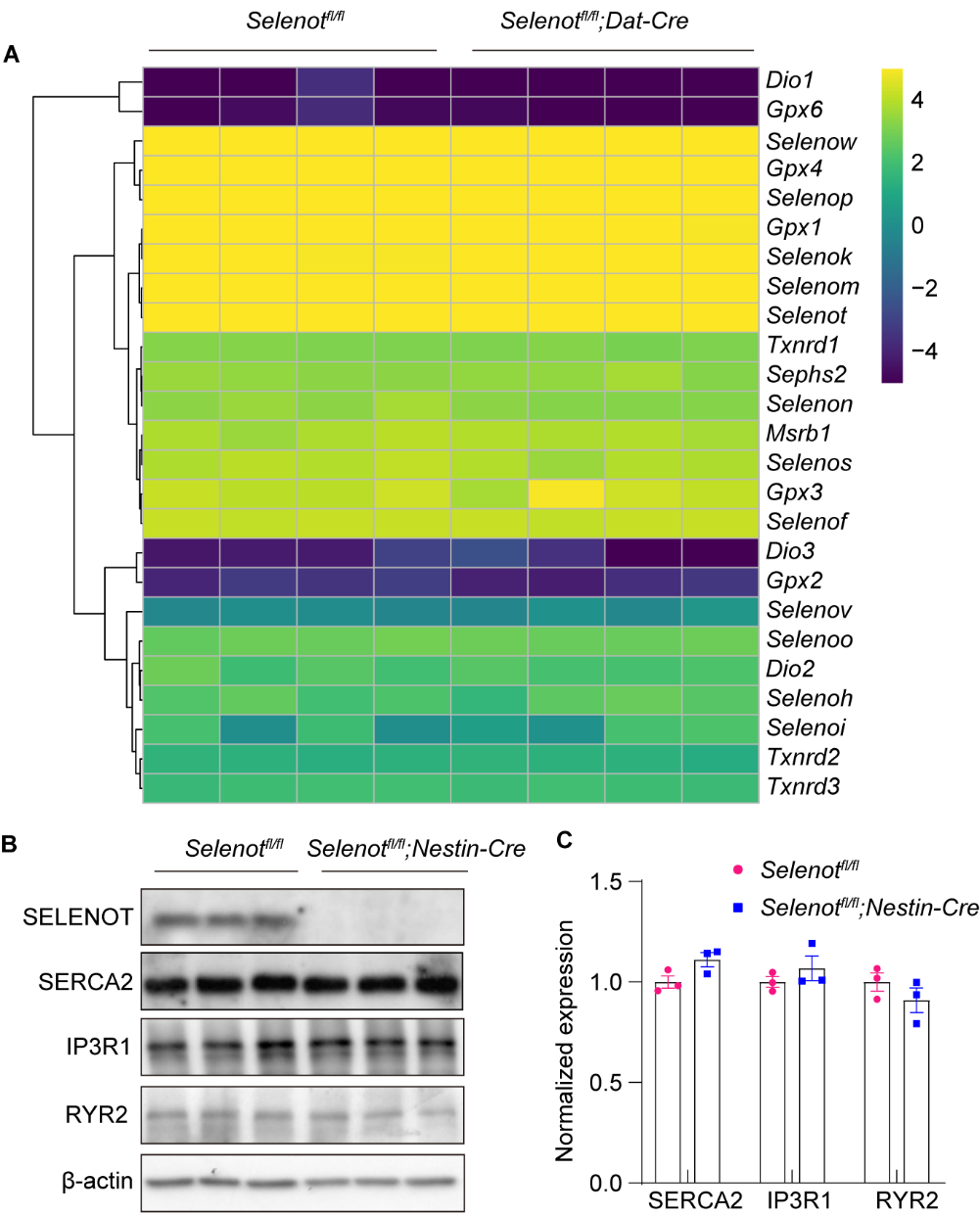

**Supplementary Figure 7. RNA-seq results of selenoprotein expression and Western blot analyses for SERCA2, IP3R1, and RYR2 expression.** (**A**) Row-normalized RNA-seq heatmap for 25 selenoprotein genes in the substantia nigra of *Selenot^fl/fl^* and *Selenot^fl/fl^;Dat-cre mice.* n = 4. (**B** and **C**) Western blot analyses (**C**) and quantifications (**C**) of SERCA2, ITPR1, and RYR2 in the cortex of *Selenot^fl/fl^* mice (n = 3) and *Selenot^fl/fl^;Nestin-cre* mice (n = 3). Quantifications are normalized to β-actin. Data are presented as means ± SEM and analyzed by two-tailed unpaired *t*-test. IP3R1, inositol 1,4,5-triphosphate receptor, type 1; RYR2, ryanodine receptor 2; SERCA2, sarco-ER Ca^2+^ ATPase 2.

**Supplementary Table 1. Primers and siRNA sequences**

|  | Forward primer, 5’-3’ | Reverse primer, 5’-3’ |
| --- | --- | --- |
| **Genotyping** |  |  |
| *Selenot* | TGCATTTTTCTGTCTCTGGGGAAG | CCTCCACGAACAGAGAAGTCAAAGG |
| *cre* | GACCAGGTTCGTTCACTCA | TAGCGCCGTAAATCAAT |
| **qPCR** |  |  |
| *DAT* (Human) | TTCTCCTGTCCGTCATTGGC | CGTGAAGCCCACACCTTTCA |
| *Dat* (Mouse) | TTCATGGTTATTGCCGGGATG | TGTAGAAGAAGCCCACGTAGAA |
| *ACTB* (Human) | TGGCACCCAGCACAATGAA | CTAAGTCATAGTCCGCCTAGAAGCA |
| *Actb* (Mouse) | GCAGGAGTACGATGAGTCCG | ACGCAGCTCAGTAACAGTCC |
| *NURR1* (Human) | ACCACTCTTCGGGAGAATACA | GGCATTTGGTACAAGCAAGGT |
| *Nurr1* (Mouse) | CAGCTCCGATTTCTTAACTCCAG | AGGGGCATTTGGTACAAGCAA |
| **siRNA, 5’-3’** |  |  |
| siCtrl | UUCUCCGAACGUGUCACGUTT |  |
| siSELT-1 | GTCAGTCTTCAAACTAGTATT |  |
| siSELT-2 | GCUUCUGCUGCUUCUCCUATT |  |
| siNURR1-1 | AAGAAGTGGTTCGCACAGACA |  |
| siNURR1-2 | AGAAAUCGGAGCUGUAUUCUC |  |

*ACTB/Actb*, β-actin; *DAT*/*Dat*, dopamine transporter; *NURR1*/*Nurr1*, nuclear receptor-related 1; siCtrl, control siRNA; siNURR1, *NURR1* siRNA; siRNA, small interfering RNA; siSELT, *SELENOT* siRNA

**Supplementary Table 2. List of Antibodies**

| Antibody | Source (catalog No.) | Host | Dilution | |
| --- | --- | --- | --- | --- |
|  |  |  | WB | IF/IHC |
| **Primary antibodies** |  |  |  |  |
| β-actin | Cell Signaling (4970S) | Rabbit | 1:2000 |  |
| BCL-2 | Abmart (T40056) | Rabbit | 1:1000 |  |
| BIP | Proteintech (11587-1-AP) | Rabbit | 1:1000 |  |
| Caspase 3 | Cell Signaling (9662S) | Rabbit | 1:1000 |  |
| Caspase 3, cleaved | Cell Signaling (9664S) | Rabbit | 1:1000 |  |
| CHOP | Cell Signaling (2895T) | Mouse | 1:1000 |  |
| COMT | Abcam (ab126618) | Rabbit | 1:1000 |  |
| DAT | Millipore (MAB369) | Rat | 1:1000 |  |
| DAT | Proteintech (22524-1-AP) | Rabbit | 1:1000 |  |
| D1R | Santa Cruz Biotech. (sc-33660) | Mouse | 1:1000 |  |
| D2R | Abcam (AB5084P) | Rabbit | 1:1000 |  |
| GAPDH | Abways (AB0037) | Rabbit | 1:2000 |  |
| MAO-A | Santa Cruz Biotech. (sc-271123) | Mouse | 1:500 |  |
| MAO-B | Santa Cruz Biotech. (sc-515354) | Mouse | 1:500 |  |
| NURR1 | Proteintech (10975-2-AP) | Rabbit | 1:1000 |  |
| SELENOT | Sigma-Aldrich (HPA039780) | Rabbit | 1:200 |  |
| SELENOT | Novus Biological (NBPI-90979) | Rabbit |  | 1:200 |
| TH | Proteintech (25859-1-AP) | Rabbit |  | 1:1000 |
| TH | Immunostar (22941) | Mouse |  | 1:1000 |
| TH | Santa Cruz Biotech. (sc-25269) | Mouse | 1:1000 |  |
| VMAT2 | Proteintech (20873-1-A) | Rabbit | 1:1000 |  |
| SERCA2 | Thermo Fisher (MA3-919) | Mouse | 1:1000 |  |
| IP3R1 | Proteintech (19962-1-AP) | Rabbit | 1:1000 |  |
| RYR2 | Proteintech (19765-1-AP) | Rabbit | 1:1000 |  |
| c-MYC | Proteintech (10828-1-AP) | Rabbit | 1:1000 |  |
| c-MYC | Proteintech (67447-1-Ig) | Mouse | 1:1000 |  |
| **Secondary antibodies** |  |  |  |  |
| Anti-mouse | Thermo Fisher (7076) |  | 1:2000 |  |
| Anti-rabbit | Thermo Fisher (7074) |  | 1:2000 |  |
| Alexa Fluor 555-conjugated anti-rabbit IgG | Thermo Fisher (A-21429) |  |  | 1:2000 |
| Alexa Fluor 488-conjugated anti-mouse IgG | Thermo Fisher (A11001) |  |  | 1:2000 |

BCL-2, B-cell lymphoma-2; BIP, binding-immunoglobulin protein; CHOP, CCAAT-enhancer-binding protein homologous protein; COMT, catechol-O-methyltransferase; DAT, dopamine transporter; D1R, dopamine D1 receptor; D2R, dopamine D2 receptor; NURR1, nuclear receptor-related 1; IF/IHC, immunofluorescence/immunohistochemistry; IP3R1, inositol 1,4,5-triphosphate receptor type 1; MAO-A, monoamine oxidase A; MAO-B, monoamine oxidase B; RYR2, ryanodine receptor 2; SELENOT, selenoprotein T; SERCA2, sarco-ER Ca^2+^ ATPase; TH, tyrosine hydroxylase; VMAT2, vesicular monoamine transporter 2; WB, Western blot.
